# VRN2-PRC2 facilitates light-triggered repression of PIF signaling to coordinate growth in Arabidopsis

**DOI:** 10.1101/2024.04.22.590552

**Authors:** Rory Osborne, Anne-Marie Labandera, Alex J. Ryder, Anastasia Kanali, Oluwatunmise Akintewe, Maximillian A. Schwarze, Christian D. Morgan, Tianyuan Xu, Sjon Hartman, Eirini Kaiserli, Daniel J. Gibbs

**Affiliations:** School of Biosciences, University of Birmingham, Edgbaston, B15 2TT, UK; School of Molecular Biosciences, College of Medical, Veterinary and Life Sciences, University of Glasgow, Glasgow, G12 8QQ, UK; Plant Environmental Signalling and Development, Faculty of Biology, University of Freiburg, Freiburg 79104, Germany; CIBSS-Centre for Integrative Biological Signalling Studies, University of Freiburg, Freiburg 79104, Germany; Université de Bordeaux, UMR Ecophysiologie et Génomique Fonctionnelle de la Vigne, 210 Chemin de Leysotte, 33140, Villenave d’Ornon, France

## Abstract

The polycomb protein VERNALIZATION2 (VRN2) is a plant-specific subunit of the polycomb repressive complex 2 (PRC2), a conserved eukaryotic holoenzyme that represses gene expression by depositing the histone H3K27me3 mark in chromatin. Previous work established VRN2 as an oxygen-regulated target of the N-degron pathway that may function as a sensor subunit connecting PRC2 activity to the perception of positional and environmental cues. Here we show that VRN2 is enriched in hypoxic meristematic regions and emerging leaves of *Arabidopsis* under non-stressed conditions, and that *vrn*2 mutants are larger than wild type, indicating that VRN2-PRC2 negatively regulates growth and development. This growth phenotype is caused by ectopic expression of genes that promote cell expansion, including many *SAUR* genes and other direct PIF transcription factor targets. Analysis of *SAUR19* promoter activity and expression dynamics revealed that VRN2 is required to specifically repress these genes in the light. Moreover, we show that VRN2 is epistatic to PIF4, and directly binds and methylates histones of key loci in the PIF4 transcriptional network to provide robust light-responsive control of gene expression and growth. We propose that hypoxia-stabilised VRN2-PRC2 sets a conditionally repressed chromatin state at PIF-regulated hub genes early in leaf ontogeny coinciding with the cell division phase, and that this is required for enhancing their subsequent repression via a light-responsive signalling cascade as cells enter the expansion phase. Thus, we have identified VRN2-PRC2 as core component of a spatially regulated and developmentally encoded epigenetic mechanism that co-ordinates environment-responsive growth by facilitating light-triggered suppression of PIF signalling.

## Introduction

Changes in chromatin structure play an important role in regulating gene expression in eukaryotes. For example, a diverse range of modifications on histone tails - including methylation (me), acetylation (ac) and monoubiquitylation - can influence gene expression though altering histone compaction, DNA accessibility, and transcription factor occupancy (Bannister and Kouzarides, 2011). One well characterised histone mark is Histone H3 Lysine 27 trimethylation (H3K27me3), a repressive modification that is deposited by the polycomb repressive complex 2 (PRC2) in most animals and land plants (Margueron and Reinberg, 2011, Simon and Kingston, 2009). Like other epigenetic histone marks, H3K27me3 is often mitotically stable, potentiated by a PRC2-mediated “read and write” mechanism that transmits the modification as DNA replicates to trigger long-term gene repression (Yu et al., 2019). However, H3K27me3 can also be erased by specific demethylases, or antagonised by histone acetyltransferases (HATs) that deposit the active H3K27ac mark (Pasini et al., 2010, Dimitrova et al., 2015, Crevillen, 2020). Thus, PRC2-mediated H3K27me3 typically functions as part of broader set of chemical changes that dynamically control the local chromatin state to fine tune promoter accessibility and consequent gene expression.

Initially discovered in *Drosophila*, the PRC2 is conserved throughout multicellular eukaryotes, and has been broadly implicated in the control of cell identity, developmental transitions and the establishment of environmental memory (Vijayanathan et al., 2022). The canonical complex is a holoenzyme containing four key subunits, and typically acts in association with an array of accessory proteins and non-coding RNAs that help to specify its functions and gene targets. In contrast to animals, several core subunits of the PRC2 have expanded in number in plants, indicating flexibility in the composition and activity of individual complexes in this kingdom (Derkacheva and Hennig, 2014, Hennig and Derkacheva, 2009, Bieluszewski et al., 2021). In *Arabidopsis thaliana* (hereafter Arabidopsis) for example, PRC2s are classified according to which of three homologs of the *Drosophila* SUPRESSOR OF ZESTE 12 (Suz12) subunit they recruit (**Fig. 1a**). This includes the sporophyte-specific Suz12 orthologs EMBRYONIC FLOWER 2 (EMF2) and VERNALIZATION 2 (VRN2), which can interchangeably associate with the two sporophyte methyltransferases CURLY LEAF (CLF) and SWINGER (SWN), as well as gametophyte-specific FERTILIZATION INDEPENDENT SEED 2 (FIS2) and its cognate methyltransferase MEDEA (MED). FERTILIZATION INDEPENDENT ENDOSPERM (FIE) is the sole homolog of the Drosophila EXTRA SEX COMBS (ESC) subunit, and although five putative variants of *Drosophila* p55 are present in the Arabidopsis genome, MULTIPLE SUPPRESSOR OF IRA 1 (MSI1) is the predominant PRC2-associated isoform. Several discrete and overlapping functions for each of these different Arabidopsis complexes have been ascribed (Derkacheva and Hennig, 2014, Hennig and Derkacheva, 2009, Margueron and Reinberg, 2011, Mozgova and Hennig, 2015, Bieluszewski et al., 2021).

**Figure 1:**
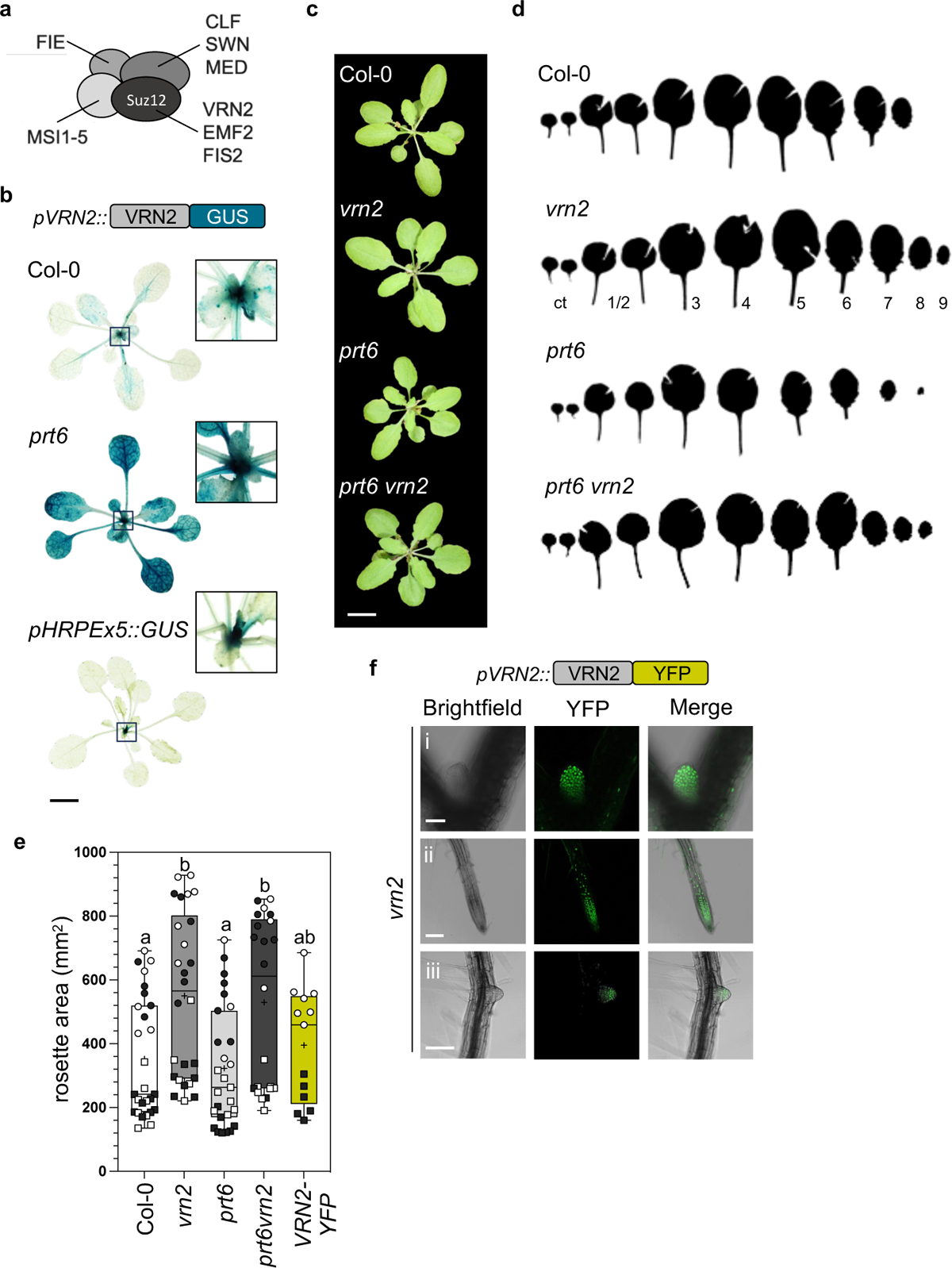
VRN2 represses rosette growth. (**a**) Schematic showing *Arabidopsis thaliana* proteins making up the four core subunits of the PRC2. VRN2 is one of three Arabidopsis homologs of Drosophila Suz12. (**b**) Histochemical staining of 4-week-old Col-0 (top) and *prt6-1* (middle) rosettes expressing the same *pVRN2::VRN2-GUS* transgene, and Col-0 expressing *pHRPEx5::GUS* (bottom). scale bar, 1cm. (**c**) Rosettes of 3-week-old Col-0, *vrn2-5*, *prt6-1* and *prt6-1vrn2-5* lines grown under long day conditions. Images were digitally extracted for comparison. Scale bar, 1cm. (**d**) Leaf series from an individual plant of Col-0, *vrn2-5*, *prt6-1* and *prt6-1vrn2-5* lines grown as in (**c**). Cotyledon (ct) and leaf number (1-9) is shown under *vrn2-5* (leaf 1 and 2 emerge simultaneously and cannot be differentiated). Cuts were made in the leaf blades to flatten them for area measurements**. (e)** Total rosette area of Col-0, *vrn2-5*, *prt6-1*, *prt6-1vrn2-5 and vrn2-5/VRN2-YFP* lines. Black and white squares and circles correspond to 4 independent biological repeats. Plot shows maximum and minimum, 25^th^ to 75^th^ percentiles, median (horizontal line) and mean (+). Statistically significant differences are shown with letters and were calculated by one-way ANOVA followed by Tukey’s test (*p*<0.05). (**f**) Confocal images of (i) SAM (ii) RM and (iii) stage VIII LRP in 7-day-old *vrn2-5/VRN2-YFP* seedling. Scale bars: (i) 75 μm, (ii, iii) 150 μm.

For example, FIS2-PRC2 prevents seed development prior to fertilization, whilst EMF2 suppresses flowering to maintain vegetative growth (Luo et al., 1999, Yoshida et al., 2001). Interestingly, whilst mutations in most of the core Arabidopsis PRC2 components, including FIS2 and EMF2, lead to pleiotropic and often severe developmental defects (Derkacheva and Hennig, 2014), this is not the case for VRN2, suggesting that VRN2-PRC2 has more specific or specialised roles. Indeed, VRN2-PRC2 has been linked to a number of environmental and developmental processes, including the regulation of vernalization and flowering time (Gendall et al., 2001, Labandera et al., 2021), seed development and dormancy (Auge et al., 2017, Roszak and Kohler, 2011), preventing somatic cell de-differentiation (Ikeuchi et al., 2015), and repressing root development (Labandera et al., 2021). Apart from vernalization, where VRN2-PRC2 is part of the mechanism that epigenetically silences *FLOWERING LOCUS C (FLC)* (Gendall et al., 2001, Yang et al., 2017), the specific targets of VRN2 involved in many of these responses are unknown or only partially characterised. VRN2-PRC2 can also associate with several plant-specific accessory proteins - including the cold-induced VERNALIZATION INSENSITIVE 3 (VIN3) and its constitutively expressed homolog VIN3-LIKE 1(VIL1)/VERNALIZATION5 (VRN5) - that help to specify its targeting and activity across the genome (Kim et al., 2021, Sung and Amasino, 2004, Kim et al., 2023, Fiedler et al., 2022, Franco-Echevarria et al., 2023).

There is increasing evidence that tissue-specific expression and posttranslational control of PRC2 subunits contribute to spatiotemporal and signal-triggered functions for the complex (Bieluszewski et al., 2021, de Lucas et al., 2016). For example, VRN2 protein stability is controlled by the PROTEOLYSIS6 (PRT6) N-degron pathway of protein degradation, mediated by a specific N-terminal (Nt-) sequence in VRN2 that is conserved and functional in VRN2-like homologs throughout the flowering plant lineage, but absent from other Suz12 homologs in plants and animals (Gibbs et al., 2018, Labandera et al., 2021, Holdsworth and Gibbs, 2020, Zhang et al., 2024). In Arabidopsis, *VRN2* mRNA is broadly expressed, but once translated the VRN2 protein is subjected to a series of oxygen (O_2_) and NO-dependent enzymatic Nt-processing events that catalyse its degradation by the 26S proteasome. As such, VRN2 is highly labile, but is relatively enriched in hypoxic niches of the plant (Weits et al., 2021) - namely the shoot apical meristem (SAM), root meristem (RM) and lateral root primordia (LRP). We previously showed that VRN2 negatively regulates root growth in Arabidopsis, suggesting it has a repressive role in hypoxic niches of the root (Labandera et al., 2021). Moreover, VRN2 is stabilised throughout plant tissues in response to environmental signals that inhibit its turnover, such as cold-exposure and flooding-induced hypoxia, and *vrn2* mutants are sensitive to low-O_2_ stress and root waterlogging (Wood et al., 2006, Gibbs et al., 2018). This suggests that VRN2 may act as a “sensor” subunit of the PRC2, allowing plants to co-ordinate H3K27me3 deposition in response to positional and environmental cues that influence O_2_ and NO availability, or otherwise inhibit the N-degron pathway.

Here we sought to investigate broader roles for VRN2 in the control of plant development, with a view to understanding its function in aerial tissues where it accumulates post-translationally under non-stressed conditions. We found that VRN2 is enriched in the meristem and emerging leaves of WT plants, and that *vrn2* mutants have larger rosettes due to increased cell size. Using a combination of genetic, molecular biology, RNA-seq and ChIP-seq approaches we show that VRN2-PRC2 negatively regulates plant growth by repressing the PIF4 regulated transcriptome – including a large number of auxin signalling and response genes - specifically in the light. A spatiotemporal analysis of VRN2 protein abundance and localisation suggests that VRN2-PRC2 functions early on in leaf organogenesis, facilitating the deposition of H3K27me3 in specific growth promoting genes in dividing cells of the hypoxic shoot meristem and young leaves. We propose that this function is important for setting a conditionally repressed epigenetic state at these loci, allowing for subsequent light-mediated inhibition of PIF signalling in the cell-expansion phase of leaf development. As such, we identify the VRN2-PRC2 module as an important epigenetic regulator of light-responsive growth and development in Arabidopsis.

## Results

### VRN2 represses rosette growth

In WT (Col-0) Arabidopsis rosettes, VRN2 protein (VRN2-GUS) is enriched in the shoot apical meristem (SAM) and emerging leaf primordia, with further weak signal in the petioles and parts of the vasculature (**Fig. 1b**). This pattern is similar to that seen for *pHRPEx5::GUS*, a hypoxia reporter that is also enriched in the SAM and leaf primordia (Weits et al., 2019). This localisation pattern is the result of post-translational degradation of VRN2 via the N-degron pathway, since VRN2-GUS accumulates to a high level in all tissues in *prt6*-*1* (**Fig. 1b**). We observed that *vrn2-5* and *prt6-1vrn2-5* mutant rosettes were larger than Col-0 and *prt6-1*, respectively (**Fig. 1c**). To quantify this, we dissected and measured the area of individual cotyledons and leaves, which showed that the increased rosette size is due to consistently larger leaves across their developmental chronology (**Figs. 1d, e and Fig S1a**). Correlating with this, *vrn2-5* rosettes had significantly increased aerial fresh weight compared to WT (**Fig. S1b**). To test that this growth phenotype was a consequence of loss of VRN2 function, we generated a transgenic *vrn2-5* line expressing a VRN2-YFP fusion driven by ∼2Kb of the native *VRN2* promoter (hereafter: *vrn2-5/VRN2-YFP*), and confirmed its nuclear localisation, and characteristic posttranslational enrichment in hypoxic SAM, RM and LRP regions (**Fig. 1f and Figs S1c-d**) (Gibbs et al., 2018, Labandera et al., 2021). This transgene reverted rosette size and rosette fresh weight to WT levels (**Fig 1e and Fig S1b, e-g**), indicating that the *VRN2-YFP* fusion protein is functional and sufficient to complement the ectopic growth phenotype of *vrn2-5*.

### Upregulation of growth promoting genes in *vrn2* seedlings and rosettes

We previously showed that *vrn2-5* mutant seedlings have significantly longer primary roots and increased lateral root density relative to WT plants (Labandera et al., 2021). Together with our observations in rosettes (**Fig. 1**), this suggests that VRN2 represses growth at different stages of the plant lifecycle. To identify VRN2 target genes responsible for these phenotypes, we conducted an RNA sequencing (RNA-seq) analysis on several mutants at different developmental stages: (1) Col-0, *vrn2-5*, *prt6-1*, and *prt6-1vrn2*-*5* at the 11-day-old seedling stage, and (2) Col-0 and *vrn2-5* at the 3-week-old rosette stage (**Fig. 2a**). Given that VRN2-PRC2 represses gene expression, we specifically looked for genes significantly upregulated (>1.5-fold, *p*<0.05) in *vrn2-5* lines vs respective controls. Here, we identified 139 upregulated genes in *vrn2-5* vs Col-0 seedlings, 104 upregulated genes in *prt6-1vrn2-5* vs *prt6-1* seedlings, and 300 genes upregulated in *vrn2-5* vs Col-0 rosettes (**Fig. 2a**). By filtering for genes that were upregulated across at least two or all three comparisons, we identified a core set of 45 genes that were significantly enriched for GO terms including “*response to auxin*”, “*response to red or far-red light”*, “*shade avoidance”*, and “*growth*” (**Figs 2a and b**). In contrast there was almost no overlap in the down-regulated gene sets, where *VRN2* was the only gene common to all three comparisons (**Fig. S2a**).

**Figure 2:**
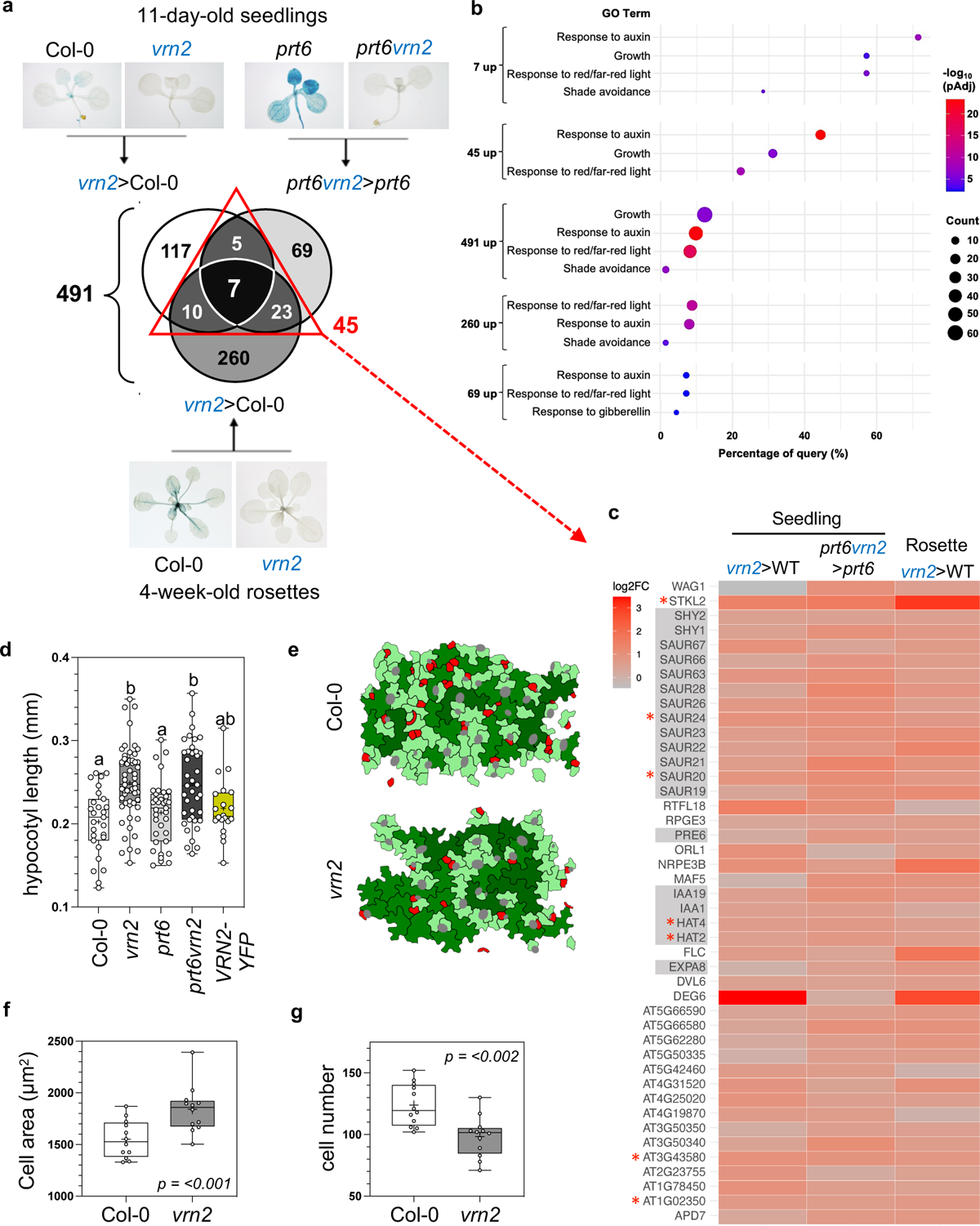
Upregulation of growth promoting genes in *vrn2* seedlings and rosettes. (**a**) Schematic and Venn diagram describing the RNA sequencing strategy and results in seedlings (top) and rosettes (bottom). 491 *vrn2-5* upregulated genes were identified across the three comparisons, of which 45 (red triangle) were found to be present in at least two comparisons. Histochemically stained plants indicate localisation of VRN2-GUS in respective mutants for descriptive purposes. (**b**) Dot plot showing the GO term enrichment of respective subsets of genes identified from the Venn diagram in (a). The Y-axis indicates the term, X-axis indicates the percentage of positive terms present for each query, dot size indicates number of genes enriched in term, and colour indicates the - log_10_ of the adjusted p-value. (**c**) Heatmap showing the log_2_ fold change in respective comparisons of the 45 *vrn2-5* UP genes denoted by the red triangle in (a). Red asterisks indicate the 7 genes that were upregulated in all 3 comparisons; grey boxes indicate genes associated with the GO terms ‘*response to auxin*’ and ‘*response to red or far-red light*’ as described in (b). (**d**) Box and whisker plot quantifying the hypocotyl lengths of 7-day old light grown seedlings of Col-0, *vrn2-5*, *prt6-1, prt6-1vrn2-5* and *vrn2-5*/*VRN2-YFP*. Data comprise 3 biological replicates. Letters indicate statistical differences calculated using 1-way ANOVA followed by Tukey’s test. (**e**) Representative images of epidermal cells from Col-0 and *vrn2-5*. Cells were analysed in ImageJ and are coloured according to cell area; red = 85-300 µm^2^, light green = 300-1600 µm^2^, mid green = 1600-3200 µm^2^, dark green >3200 µm^2^. Grey cells are guard cells/stomata. (**f**) Box and whisker plot quantifying the average area of epidermal cells from leaf 3 of 2 week old Col-0 and *vrn2-5* plants. Plot shows maximum and minimum, 25^th^ to 75^th^ percentiles, median (horizontal line) and mean (+). Statistical difference was calculated using a student’s t-test. Absolute *p-value* is displayed on the plot. Data comprise 2 images from 6 independent plants for each genotype. (**g**) Box and whisker plot quantifying the number of epidermal pavement cells per 0.24 mm^2^ from leaf 3 of 2 week old Col-0 and *vrn2-5* plants. Plot shows maximum and minimum, 25^th^ to 75^th^ percentiles, median (horizontal line) and mean (+). Statistical difference was calculated using a student’s t-test, absolute *p-value* is displayed on the plot. Data comprise 2 images from 6 independent plants for each genotype.

Amongst the 45 *vrn2-5* up DEGs were 11 *SMALL AUXIN UPREGULATED RNA* (*SAUR*) genes, several transcription factors of the *HAT/ARABIDOPSIS THALIANA HOMEOBOX PROTEIN (ATHB)* family (notably *HAT2* and *HAT4/ATHB-2*), various auxin signalling genes (*SHORT HYPOCOTYL 1 (SHY1)/INDOLE-3-ACETIC ACID INDUCIBLE 6 (IAA6)*, *SHY2/IAA3*, *IAA1* and *IAA19*), as well as genes linked to brassinosteroid signalling and cell-wall expansion (e.g. *PACLOBUTRAZOL RESITANCE 6 (PRE6)* and *EXPANSIN 8 (EXPA8)*) (**Fig. 2c**). Additionally, many other related genes – including 7 further *SAURs*, 3 more *IAAs*, 3 more *PREs*, the auxin biosynthesis gene *YUC9*, and 2 Gibberellin biosynthesis genes - were present in individual comparisons (**Fig. S2b**). Most of these genes have previously been linked to cell-expansion mediated growth, particularly within the context of hypocotyl growth in response to far-red light or shade, leaf cell expansion, and thermomorphogenesis (Franklin et al., 2011, Sun et al., 2012, Quint et al., 2023, Spartz et al., 2012, Vanhaeren et al., 2014). This prompted us to investigate the hypocotyl length and leaf cell size of photomorphogenic seedlings. We observed longer hypocotyls in *vrn2-5* and *prt6-1vrn2-5*, which could be reverted through complementation with VRN2-YFP (**Fig 2.d**), and an increase in epithelial pavement cell area in *vrn2-5 vs* WT (**Figs. 2e-g**). These findings, coupled with the observation that there was no significant difference in the dry weight of Col-0 and *vrn2-5* at the rosette stage (**Fig. S1b**), indicates that the larger size of plants lacking VRN2 can be attributed to increased cell expansion. Thus, our RNA sequencing analysis identified a broad set of upregulated genes in *vrn2-5* seedings and adult plants that are known to promote growth.

### VRN2 is required for spatiotemporal and light-triggered repression of *SAURs*

SAURs are a large group of plant-specific proteins that promote cell expansion by inhibiting PP2C-D phosphatases, which in turn activate plasma membrane H^+^-ATPases to drive acid growth (Ren and Gray, 2015, Stortenbeker and Bemer, 2019). Given the large number of *SAURs* present in the shared *vrn2-5* up DEGs (11/45; **Fig. 2c**), we investigated the connection between VRN2 and *SAUR* expression further. We focussed on *SAUR19*, the archetypal member of the *SAUR19-24* sub family that is fully represented in the RNA-seq. We crossed a *pSAUR19::GUS* reporter line (Spartz et al., 2012) to *vrn2-5*, *prt6-1* and *prt6-1vrn2-5* and monitored promoter activity by anti-GUS Western blotting (**Figs. 3a and S3a**). When comparing GUS signal in skotomorphogenic seedlings, no difference in promoter activity was observed. However, in photomorphogenic seedlings and rosette leaves harvested in the light, GUS levels were much higher in *vrn2-5* and *prt6-1vrn2-5* compared to Col-0 and *prt6-1*, indicating elevated *SAUR19* promoter activity, in support of our RNA-seq data. *SAUR19* is highly expressed in dark grown seedlings and rapidly downregulated during de-etiolation (Sun et al., 2016). Directly comparing *pSAUR19::GUS* activity in etiolated vs photomorphogenic seedlings on the same blot confirmed these dynamics in Col-0, but showed that this repression is compromised in *vrn2-5* (**Fig. S3b**). In addition, a qRT-PCR analysis of *SAUR19* and *SAUR24* expression over a diurnal time course revealed similar mRNA levels in Col-0 and *vrn2-5* at ZT 22 and ZT 0 (i.e., near the end of the night and dawn, respectively), but a much stronger induction of expression at ZT4 and ZT8 in *vrn2-5* relative to Col-0 (**Figs. 3b and S3c**). This corroborated the *pSAUR19::GUS* data, showing that ectopic expression of *SAURs* in *vrn2-5* is light-responsive rather than constitutive. We also examined the spatial pattern of ectopic *pSAUR19::GUS* expression by histochemical staining in 7- and 10-day-old seedlings and 4-week-old rosettes (**Fig. 3c and S3d**). In Col-0, *pSAUR19::GUS* expression is largely restricted to hypocotyl, petiole, and vasculature regions, but in *vrn2-5* it is expressed much more broadly throughout the whole plant, including the expansion zone of leaves. To investigate the temporal dynamics of this misexpression further, we stained seedlings at 1, 3, 5 and 7 days after germination. Spatial differences in *GUS* expression were only observed from day 5, indicating that ectopic *pSAUR19* activity in *vrn2-5* is initiated during seedling establishment (**Fig. S3e**). Thus, our data reveal that VRN2 is required for light-triggered and spatiotemporal repression of *SAURs*.

**Figure 3.**
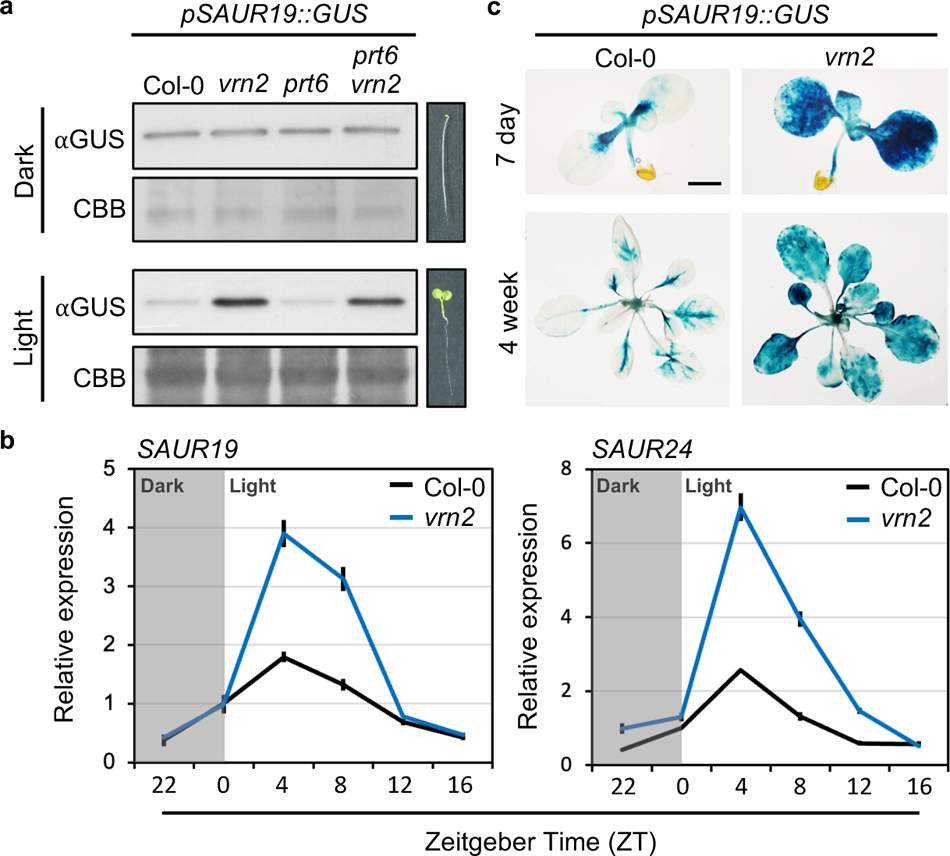
VRN2 is required for spatiotemporal and light-triggered repression of *SAURs*: (**a**) anti-GUS Western blot of *pSAUR19::GUS* in Col-0, *vrn2-5*, *prt6-1* and *prt6-1vrn2-5* in dark (top) and light (bottom) grown seedlings. Coomassie brilliant blue (CBB) staining was used to quantify equal loading. Seedling images are for descriptive purposes only. GUS band is ∼68 kDa. (**b**) qRT-PCR quantifying *SAUR19* and *SAUR24* expression in Col-0 and *vrn2-5* at the indicated timepoints displayed in Zeitgeber (ZT) time. Data were normalised to 2 housekeeping genes and are presented relative to Col-0 at ZT 0. Each point represents the mean of 2 technical repeats from one biological replicate; error bars represent the standard error of the mean. 3 independent biological replicates showing a similar trend can be found in Fig S3. (**c**) Histochemical staining of representative Col-0 and *vrn2-5* plants expressing *pSAUR19::GUS* at the 7 days (top) and 4 weeks (bottom) of growth. Scale bars = 1 mm (top) and 1 cm (bottom) respectively.

### The VRN2-PRC2 module is epistatic to PIF4

The light-dependent nature of VRN2-mediated *SAUR19* repression (**Fig. 3**), coupled with the enrichment of genes linked to *response to red/far-red light* GO terms in the *vrn2* up DEGs (**Fig. 2b**) led us to investigate VRN2 function within the context of phytochrome (Phy) signalling. Phytochromes are the primary red/far red-light receptors in plants and transduce light signals into transcriptional changes through degrading and inhibiting the activity of several PHYTOCHROME INTERACTING FACTOR (PIF) transcription factors (Castillon et al., 2007, Cheng et al., 2021). We compared the *vrn2* up DEGs with several published PIF ChIP data sets and found that 71% (32/45, *p*<0.001), 53% (24/45 *p*<0.001) and 58% (26/45, *p*<0.001) are direct binding targets of PIF4, PIF5 and PIF7, respectively (**Fig. S4a**) (Oh et al., 2012, Hornitschek et al., 2012, Yang et al., 2021). Amongst the 32 reported PIF4 targets, 25 also had significantly reduced mRNA abundance in the *pif4* mutant vs WT (Gangappa et al., 2017), including all 6 members of the *SAUR19* subfamily, *HAT2*, *HAT4* and several I*AAs* (**Fig. 4a**). Due to this overlap, we chose to further explore the genetic and functional interactions between VRN2 and PIF4, by generating and phenotypically characterising a *pif4-2vrn2*-5 double mutant. The large rosette phenotype of *vrn2-5* was abolished in the *pif4-2vrn2-*5 double mutant, which was similar to the *pif4-2* single mutant (**Fig. 4b and c**). Moreover, the longer hypocotyl observed in light grown *vrn2-5* seedlings was also reverted in *pif4-2vrn2-5* (**Fig. 4d**), and this corresponded with a concomitant reduction of the elevated *SAUR19* expression to levels seen in *pif4-2* (**Figs. 4e**). Thus, mutations in *PIF4* can suppress the *vrn2-5* growth and ectopic *SAUR19* expression phenotypes, indicating epistasis and a functional connection between these loci.

**Figure 4.**
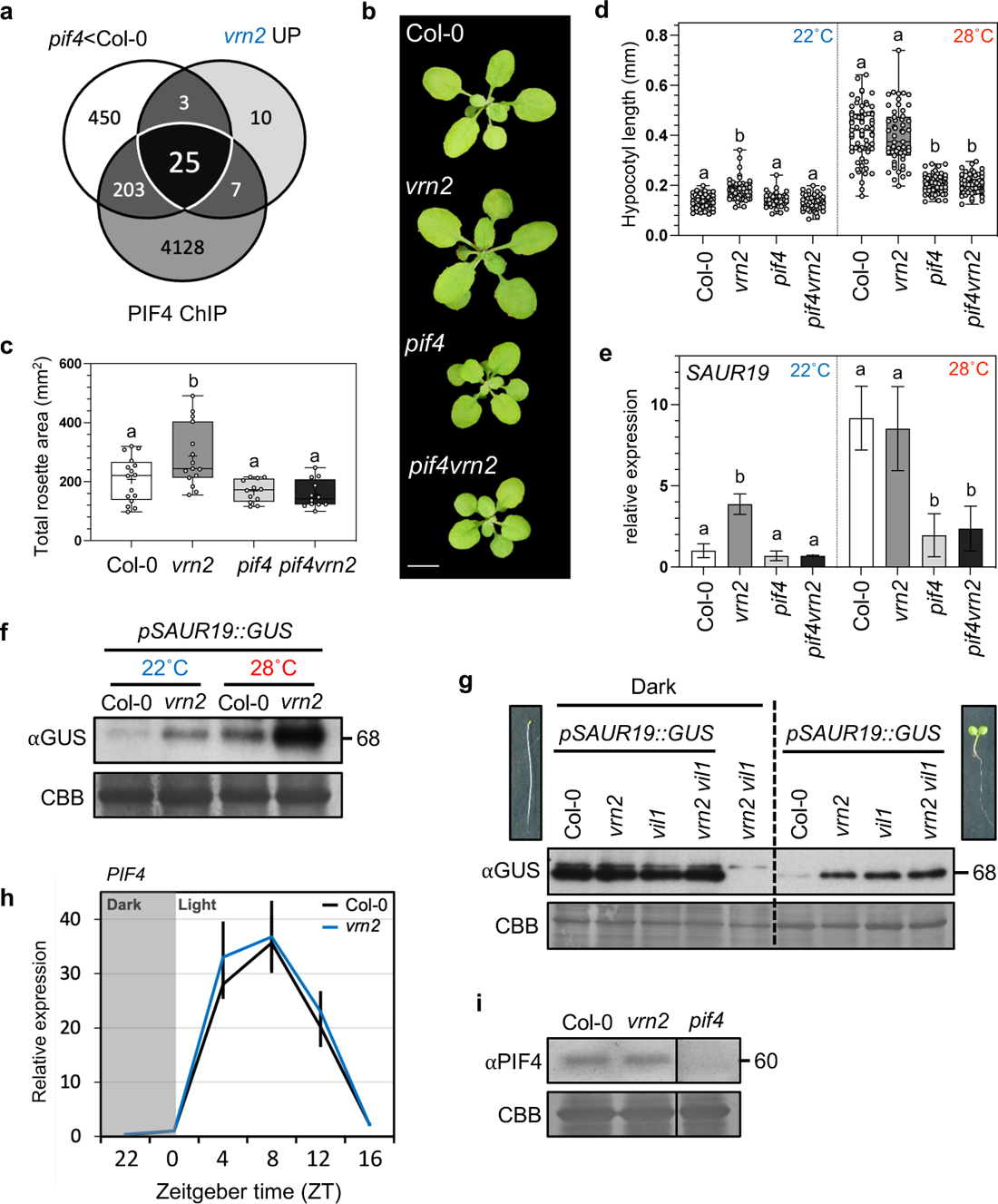
The VRN2-PRC2 module is epistatic to PIF4. (**a**) Venn diagram showing the overlap between the 45 *vrn2-5* UP genes (Fig. 2A), *pif4* down genes (Gangappa et al., 2017) and genes identified as high confidence PIF4 binding targets (Oh et al., 2013). (**b**) Representative images of 3-week-old Col-0, *vrn2-5*, *pif4-2* and *pif4-2vrn2-5* rosettes. Scale bar = 1 cm.(**c**) Quantification of total rosette area after leaf dissection of 3-week-old Col-0, *vrn2-5*, *pif4-2* and *pif4-2vrn2-5* plants. Plot shows maximum and minimum, 25^th^ to 75^th^ percentiles, median (horizontal line) and mean (+). Statistically significant differences are shown with letters and were calculated by one-way ANOVA followed by Tukey’s test (*p*<0.05). (**d**) Box and whisker plots showing hypocotyl lengths of Col-0, *vrn2-5*, *pif4-2* and *pif4-2vrn2-5* seedlings grown vertically on ½ MS agar plates for 7 days at either 22°C or 28°C. Each point represents a single seedling; data shown are pooled from 3 independent biological replicates. Letters indicate statistically significant differences calculated by 1-way ANOVA followed by Tukey’s test. (**e**) qRT-PCR quantifying *SAUR19* expression in 7-day-old seedlings of Col-0, *vrn2-5*, *pif4-2* and *pif4-2vrn2-5* at 22°C or following transfer to 28°C for 6h. Each bar represents the average of data from 2 biological replicates; error bars represent the standard error of the mean. Data were normalised to 2 housekeeping genes and are presented as relative to Col-0 at 22°C. Letters indicate statistically significant differences calculated by 1-way ANOVA followed by Tukey’s test. (**f**) Western blot showing *pSAUR19::GUS* expression in 7-day-old seedlings of Col-0 and *vrn2-5* grown at 22°C or 28°C. Coomassie brilliant blue (CBB) staining was used to quantify equal loading. (**g**) Western blot showing *pSAUR19::GUS* expression in 4-day-old seedlings of Col-0, *vrn2-5*, *vil1-1* and *vrn2-5 vil1-1* grown in the dark (left) and in the light (right). *vrn2-5 vil1-1* without *pSAUR19::GUS* was used as a negative control. Coomassie brilliant blue (CBB) staining was used to quantify equal loading between samples and growth conditions. Seedling images are for descriptive purposes only. (**h**) qRT-PCR quantifying *PIF4* expression in Col-0 and *vrn2-5* at the indicated timepoints displayed in Zeitgeber (ZT) time. Data were normalised to 2 housekeeping genes and are presented relative to Col-0 at ZT 0. Each bar represents the average of data from 3 biological replicates. (**i**) Western blot showing PIF4 protein levels at ZT4 in 7-day old seedlings of Col-0 and *vrn2-5* grown in the light. *pif4-2* protein extract was used as a negative control. Coomassie brilliant blue (CBB) staining was used to quantify equal loading between samples.

As well as coordinating light triggered growth, PIF4 is also a major regulator of thermomorphogenesis, where it promotes hypocotyl elongation in response to warm temperature via induction of auxin and brassinosteroid synthesis and signalling that ultimately converge on *SAUR* expression (Franklin et al., 2011, Sun et al., 2012, Oh et al., 2012). Hypocotyl extension in photomorphogenic seedlings grown at 28°C was significantly induced in both Col-0 and *vrn2-5* relative to 22°C, but the response was severely dampened in *pif4-2* and *pif4-2vrn2-5* double mutants (**Fig. 4d**). Final hypocotyl lengths were similar for Col-0 and *vrn2-5*, indicating a greater relative growth in Col-0 than *vrn2-5* at warm temperatures. *SAUR19* mRNA levels were similarly induced after 6h transfer to 28°C in Col-0 and *vrn2-5* (**Fig. 4e**), but restricted in *pif4-2* and *pif4-2vrn2-5*, in line with previous studies demonstrating the requirement of PIF4 in this transcriptional cascade (Franklin et al., 2011). In contrast, when we examined promoter activity in the *pSAUR19::GUS* lines grown in constant temperature, induction of GUS was proportionally greater in *vrn2-5* vs Col-0 at both 22°C and 28°C (**Fig. 4f**), suggesting that the enhanced competency for expression in *vrn2-5* vs Col-0 is maintained even at warmer temperatures. This discrepancy between elevated *SAUR19* promoter activity but equivalent *SAUR19* mRNA levels in *vrn2-5* vs WT at 28°C could be due to highly labile nature of *SAUR* mRNAs relative to *GUS* mRNA (Spartz et al., 2012). Nonetheless, our data reinforce the notion that VRN2 is required to repress growth promoting genes under light and temperature conditions where they should not be expressed.

It was recently shown that the VRN2-PRC2 accessory protein VIL1/VRN5 (hereafter VIL1) interacts and cooperates with phyB to regulate PIF-dependent expression dynamics at the *HAT4* promoter, by promoting the formation of a phyB-activated chromatin loop (Kim et al., 2021). Chromatin loop formation at the *HAT4* locus was also abolished in a *vrn2* mutant, corroborating the fact that VIL1 is a specific interactor of VRN2. In our study, *HAT4* was identified as one of only 7 genes that was ubiquitously upregulated across all three RNA-seq comparisons (**Fig. 2c**). To understand the relative contributions of VRN2 and VIL1 to light triggered regulation of *SAUR* expression, we crossed *vrn2-5/pSAUR19::GUS* to a *vil1-1* mutant and monitored promoter activity. Here we observed similar ectopic *pSAUR19::GUS* promoter activation in the light, which was not amplified in the *vrn2-5 vil1-1* double mutant, indicating a synergistic rather than additive effect (**Fig. 4g**). Interestingly however, when we compared our *vrn2-5* up DEGs with two previously published *vil1* mutant RNA seq data sets (Kim et al., 2023, Zong et al., 2022), we saw very little overlap (only 3 genes, and a complete absence of *SAURs* and *HAT4* etc) as well as an absence of GO terms linked *“red/far red”* or *“shade avoidance”* response (**Fig. S4c**). This suggest that control of gene expression by the VRN2-VIL1 module is highly dynamic and likely age and developmental stage regulated.

Since PIF4 is necessary for the ectopic growth phenotypes observed in *vrn2-5* mutants, we next tested to see if PIF4 itself is a target of VRN2. PIF4 mRNA expression levels were unaltered in *vrn2-5* relative to Col-0 under a diurnal time course (**Figs. 4h**). Steady-state levels of PIF4 protein at ZT4 were also equivalent in both backgrounds (**Fig. 4i**), further ruling out the hypothesis that VRN2 directly regulates *PIF4* expression. These findings suggest that VRN2 function in repressing growth is epistatic to PIF4 and is not a direct consequence of changes to its expression or translation.

### Identification of direct VRN2 gene targets

Our data indicate that VRN2 represses the PIF4-regulated transcriptome. However, whilst the RNA-seq analysis (**Fig. 2**) provides insight into the broader gene expression changes driving the growth phenotypes in *vrn2-5*, it cannot differentiate between direct and indirect (i.e., downstream) targets of VRN2. To resolve this, we performed protein chromatin sequencing (ChIP-seq) with VRN2-YFP expressing plants to identify VRN2-bound loci, and H3K27me3 ChIP-seq to identify genes that are hypomethylated in *vrn2-5* vs Col-0. After peak calling, 1,474 regions of the mappable Arabidopsis genome were found to interact with VRN2-YFP, whilst 703 regions exhibited reduced histone methylation in *vrn2-5*. We observed enrichment of the previously reported VRN2 targets *WIND1*, *WIND2* and *WIND3* in our VRN2-YFP ChIP (Ikeuchi et al., 2015), and *vrn2-5*-dependent depletion of H3K27me3 at the PRC2 regulated *ULT1* (also *LEC1*, *NIT2*, *AGL15*), demonstrating the validity and reliability of these datasets (**Fig. S5a and b**).

Comparison of the VRN2-YFP peaks and the H3K27me3 alignment data revealed a broad reduction in H3K7me3 signal for these regions in *vrn2-5* relative to Col-0 (**Fig. 5a**), despite the total levels of H3K27me3 being equivalent between genotypes that either lack or over accumulate VRN2 (**Fig 5b**). This suggests that VRN2 targets a specific subset of the genome rather than playing a fundamental role in catalysing global H3K27me3 deposition, in keeping with its highly labile nature and partial redundancy with EMF2 (Gibbs et al., 2018, Labandera et al., 2021, Derkacheva and Hennig, 2014). The 1474 VRN2-YFP peaks and 703 regions depleted for H3K27me3 corresponded to 1,265 and 699 genes, respectively. Of these, 118 genes were common to the two datasets, revealing that approximately 10% of VRN2-bound genes are also methylated by VRN2-PRC2 (**Fig. 5c**). These 118 genes were designated as “high confidence (HC) VRN2 targets”. Amongst them were the auxin biosynthesis genes *YUC8* and *YUC9*, the auxin efflux transporter *PIN-FORMED 1* (*PIN1*), and the abscisic acid (ABA) receptors *PYROBACTIN RESISTANCE 1 LIKE 2* (*PYL2*) and *PYL3* (**Fig. 5d**). Interestingly, YUC8 is a key intermediate in the PIF4 signalling cascade that drives *SAUR* expression (Sun et al., 2012), whilst YUC9 acts synergistically with YUC8 during the shade avoidance response (Muller-Moule et al., 2016) and is also upregulated in *vrn2-5* vs Col-0 (**Fig. S2b**). We designed primers overlapping the identified peaks and using ChIP-PCR confirmed that *YUC8*, *YUC9*, and *PIN1* are direct VRN2-YFP binding targets that have reduced H3K27me3 in *vrn2-5* relative to Col-0 (**Fig. 5f-g**).

**Figure 5.**
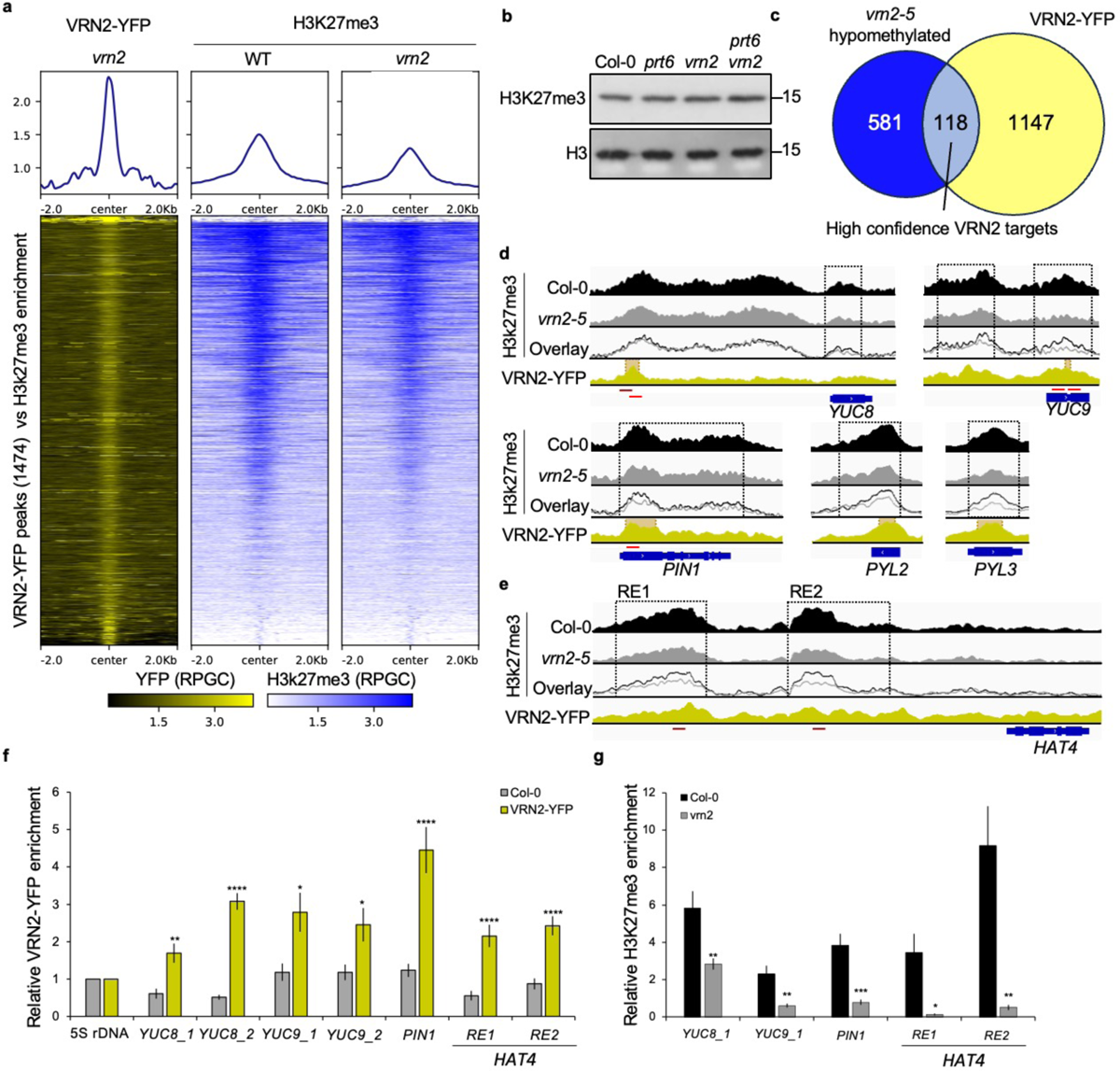
Identification of direct VRN2-PRC2 binding targets. (**a**) Coverage plots (top) and associated heatmaps (bottom) showing the distribution of H3K27me3 in Col-0 and *vrn2-5* across regions targeted by VRN2-YFP. Data show coverage across the combination of 3 biological replicates which were normalised by reads per genome coverage. (**b**) Western blot showing abundance of H3K27me3 in Col-0, *vrn2-5*, *prt6-1* and *prt6-1vrn2-5*. *⍺*H3 was used to evaluate equal loading and control for efficiency of nuclear enrichment between genotypes. Numbers indicate size (kDa). (**c**) Venn diagram showing overlap of genes (High confidence VRN2 targets) identified as depleted for H3K27me3 in *vrn2-5* (*vrn2-5* hypomethylated; blue) or targeted by VRN2-YFP (VRN2-YFP; yellow). (**d**) IGV plots showing alignment data for high confidence VRN2 targets. Black and grey tracks represent Col-0 and *vrn2-5* H3K27me3 coverage respectively; ‘overlay’ represents these same tracks mapped over one another. Black dotted boxes indicate regions which were identified as significantly depleted for H3K27me3 in *vrn2-5*. The yellow track represents VRN2-YFP coverage across the same genomic loci; orange boxes indicated regions which were identified as significantly enriched by VRN2-YFP. Data show coverage across the combination of 3 biological replicates which were normalised by reads per genome coverage. In each graph, blue tracks indicate the position and direction of the genomic sequence for each gene. (**e**) As in (d) for the *vrn2-5* hypomethylated gene *HAT4*. (**f**) ChIP-PCR of selected VRN2 targets using primers (denoted as red lines in d and e) showing enrichment in *VRN2-YFP* expressing plants. Enrichment was calculated by normalising % input values to the enrichment of *5SrDNA* for each sample. Bars represent the average value from 2 biological replicates. Error bars represent the standard error of the mean; statistically significant differences in enrichment between Col-0 and *VRN2-YFP* were calculated by students t-test (p < 0.05 *, p < 0.01 **, p < 0.0001 ****). (**g**) ChIP-PCR of HC VRN2 targets using primers (denoted as red lines in (d) and (e) showing depletion of H3K27me3 in *vrn2-5.* H3K27me3 levels were calculated by first adjusting Ct values according to the histone H3 enrichment and are presented as relative enrichment compared to the 5SrDNA negative control, for each primer pair. Bars represent the average value from 2 biological replicates. Error bars represent the standard error of the mean; statistically significant differences in enrichment between Col-0 and *vrn2* were calculated by students T-Test (p < 0.05 *, p < 0.01 **, p < 0.001 ***).

Annotated peaks identified in only one of the ChIP datasets were designated “VRN2-YFP” or “*vrn2*-hypomethylated” targets respectively (Supplementary Table S3). We posited that many of these could also represent *bona fide* VRN2-YFP targets. Amongst the latter was *HAT4* (**Fig. 5e**), which was hypomethylated at previously defined promoter response elements (REs) (Kim et al., 2021) and up-regulated in the *vrn2-5* transcriptome (**Fig 2b**). We were able to confirm VRN2-YFP binding to RE1 and RE2 in the *HAT4* promoter by ChIP-PCR, as well as reduced H3K27me3 levels in *vrn2-5* at these regions (**Fig. 5f-g**). As predicted by the Plant Chromatin State Database, despite being prevalent *vrn2-5* up DEGs, we observed neither H3K27me3 nor VRN2-YFP enrichment in *SAURs 19-24* and *60-65* (**Fig. S5c**), indicating that they are not direct methylation targets of VRN2-PRC2, and that their ectopic upregulation in *vrn2-5* mutants is a downstream consequence of loss of VRN2 function. Moreover, there is only a very small overlap (1%) between VRN2-YFP targets and *vrn2* up DEGs (**Fig. S5d**), similar to that seen when looking at PIF4 targets vs genes that are downregulated in *pif4* vs Col-0 (4.7%; **Fig. 4a**).

Overall, our combined ChIP analyses revealed a range of high confidence VRN2 target genes linked to diverse cellular processes, including flowering time, meristem identity and maintenance, and auxin signalling, and suggests that VRN2’s role in repressing PIF signalling is a consequence of it targeting a small set of “hub” genes that then trigger broader transcriptional changes to repress growth in the light.

### Spatiotemporal and light responsive analysis of VRN2 stability

VRN2-mediated repression of PIF4 signalling is light-dependent (**Figs. 3, 4**), which prompted us to explore whether this could be due to changes in *VRN2* mRNA expression and/or VRN2-YFP protein abundance across a diurnal time course (**Fig. 6a,b**). Here we saw no obvious differences in transcript levels or protein stability, suggesting that VRN2 itself is not regulated by light or time of day. We therefore considered that there might be a developmentally encoded or temporal mechanism at play, particularly because there is an obvious spatial disconnect between VRN2 accumulation in the SAM and emerging leaves, and where *SAUR19* is mis-expressed in the *vrn2-5* mutant; namely, the cell expansion zone of older leaves (**Fig. 6c**, and **Figs. 1b and 3c**). Moreover, the larger rosette size of *vrn2-5* mutants corresponds to an increase in cell size, a process that occurs in mature leaves (**Fig. 2b and S1b**). Dissected leaves of 4-week-old VRN2-GUS plants displayed a very specific pattern of VRN2-GUS localisation across leaf development (**Figs. 6d, e and Fig S6**). VRN2-GUS signal was strong throughout the meristem, leaf primordia (corresponding to leaves 11 and 10), and youngest leaf 9 (**Fig.6d**). However, VRN2-GUS signal clearly receded in a basal direction (i.e., towards the petiole) as leaves aged (**Fig. 6e**). This was restricted to the petiole in leaf 6 and was completely absent in leaves 5 to 1 (**Fig. S6**). This wave of VRN2-GUS depletion appears to correspond to the cell division arrest front, a critical zone where cells exit mitosis and subsequently contribute to leaf growth through expansion alone (Gonzalez et al., 2012, Vanhaeren et al., 2015). Moreover, this appears to be post-translationally regulated, since VRN2-GUS is broadly stable throughout all leaves in the *prt6-1* mutant, in the absence of changes in *VRN2* expression levels (**Fig. 1b**) (Gibbs et al., 2018, Labandera et al., 2021). This pattern of VRN2-GUS staining suggests that VRN2 abundance is tightly regulated across leaf development, and perhaps points to a gradient of oxygen availability as organs emerge from the hypoxic shoot apical meristem (Weits et al., 2019), divide, and then expand. As such, we propose a spatially separated and developmentally encoded mechanism whereby VRN2-PRC2 binds and methylates target genes early in leaf organogenesis when cells are dividing, establishing a mitotically stable and conditionally repressed epigenetic state that permits effective light-triggered growth-repression by the PhyB-VIL1 module (Kim et al., 2021) during cell expansion (**Fig. 6f**).

**Figure 6.**
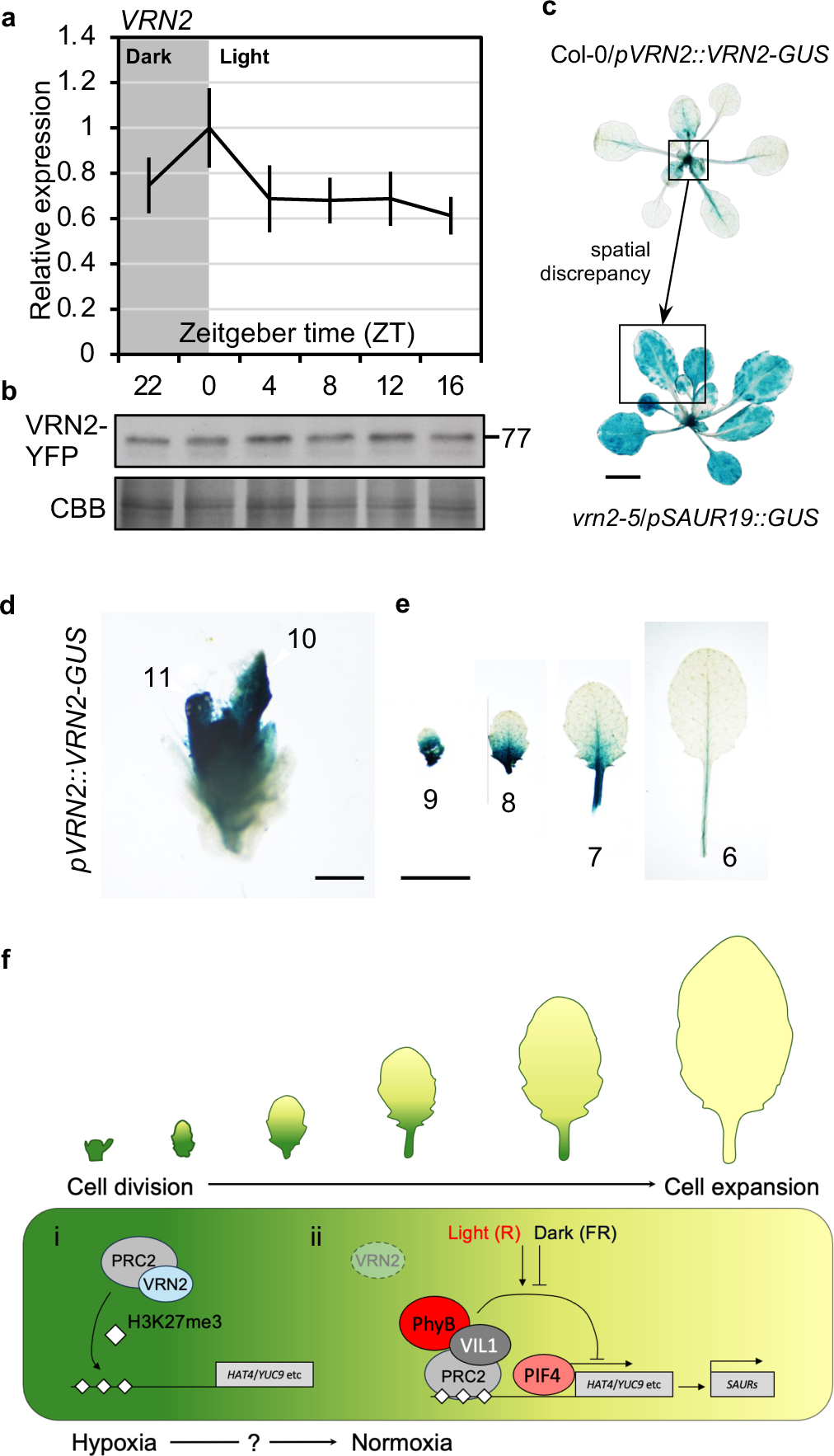
Spatiotemporal and light responsive analysis of VRN2 stability. (**a**) qRT-PCR quantifying *VRN2* expression in Col-0 at the indicated timepoints displayed in Zeitgeber (ZT) time. Data were normalised to 2 housekeeping genes and are presented relative to Col-0 at ZT 0. Each point represents the average of data from 3 biological replicates; error bars represent the SE of the mean. No significant difference was observed between any of the timepoints, as calculated by 1-way ANOVA. (**b**) Western blot showing VRN2-YFP levels in 7-day old seedlings at the indicated timepoints displayed in Zeitgeber (ZT) time. Coomassie brilliant blue (CBB) staining was used to quantify equal loading. (**c**) Histochemical staining of rosettes expressing Col-0/*pVRN2::VRN2-GUS* (top) and *vrn2-5/pSAUR19::GUS* (bottom), illustrating the spatial disconnect between where VRN2 localises and where *pSAUR19* promoter activity is ectopically induced. (**d,e**) Representative histochemical staining from a dissected 4-week-old Col-0/*pVRN2::VRN2-GUS* rosette. The rosette was dissected in developmental order to expose the meristem, leaf primordia (leaves 10 and 11) and leaves 6 – 9, all arranged in developmental order from youngest (left) to oldest (right). Scale bar in d = 500 μm. Scale bar in e = 5 mm. (**f**) Hypotheticl model of VRN2-mediated repression of growth in Arabidopsis. In the dividing cells of the hypoxic shoot meristem and emerging leaf primordia, stable VRN2 catalyses H3K27me3 deposition in promoters and gene bodies of core PIF4 activated genes (eg. *HAT4*/*YUC9*). (ii) In the expanding cells of normoxic leaf tissues, where VRN2 is degraded, this H3K27me3 mark is stable and facilitates the conditional repression of these genes in the light by the red-light activated PhyB-VIL1 module. This restricts the ectopic expression of SAURs in the cell-expansion zone in response to light. Green to yellow gradient follows the transition of division-mediated to expansion-mediated growth across developmental time as leaves emerge and grow.

## Discussion

Here we have shown that VRN2, a subunit of the PRC2, epigenetically suppresses growth in Arabidopsis through light-specific attenuation of PIF signalling, actioned through histone methylation of specific hub genes in the PIF4 transcriptional network. RNA-sequencing at both the seedling and rosette stage revealed that many *vrn2-5* upregulated genes promote cell-expansion mediated growth (**Fig. 2c and S2**), and have been previously reported as PIF4 targets, including *HAT2*, *HAT4*, *YUC9* and the *SAUR19-24* family (Franklin et al., 2011, Quint et al., 2023, Sun et al., 2012). Using a *pSAUR19::GUS* reporter, we demonstrate this gene is both upregulated and expressed ectopically in *vrn2-5* (**Figs. 3a-b and S3a-b**). Furthermore, we found that this regulation was light dependent, and only observable from 5 days after gemination (**Fig. S3e**), suggesting the regulatory role of VRN2-PRC2 is linked to the day/night cycle and is initiated during seedling establishment and not during embryogenesis. The enhanced growth phenotype in *vrn2-5* was completely abolished in the *vrn2-5pif4-2* mutant, as was the upregulation of *SAUR19* (**Figs. 4d,e**). Our data suggest that the epigenetic regulation of only a small number of genes by VRN2-PRC2 has a significant impact on the PIF-regulated transcriptional landscape. Of particular interest are *HAT4*, *YUC8* and *YUC9*, which have previously been shown to function downstream of PIF4 to activate *SAUR* expression and drive cell expansion (Muller-Moule et al., 2016, Sun et al., 2012, Franklin et al., 2011).

It was recently reported that phyB associates with the PRC2 accessory protein VIL1 to trigger the formation of a repressive chromatin loop in *HAT4,* and that this is negatively regulated through COP1-mediated degradation of VIL1 in the dark (Wang et al., 2024, Kim et al., 2021). VIL1 is a specific interactor of VRN2 (Franco-Echevarria et al., 2023), and in accordance with this, chromatin loop formation was abolished in a *vrn2* mutant (Kim et al., 2021). We observed increased *SAUR19* promoter activity in *vil1* and showed that this effect was not additive in *vrn2-5 vil1-1*, suggesting these proteins do indeed function in the same pathway to repress growth-promoting gene expression. This is supported further by our observation that VRN2-YFP physically associates with two regulatory elements critical for loop formation in the *HAT4* promoter, that also exhibit reduced H3K27me3 deposition in *vrn2-5* relative to wildtype (**Fig 5e-f**). Whether other VRN2 targets are also regulated by phyB/VIL1-dependent chromatin loop formation requires further investigation. It will also be important to determine which other PRC2 subunits interact with VRN2 as it targets these loci.

Several observations suggest that the larger phenotype of *vrn2-5* is due to increased cell expansion: (1) pavement cells of *vrn2-5* mutant are larger than in Col-0 (**Fig. 2e**); (2) *vrn2-5* mutant rosettes have increased fresh - but not dry – weight (Fig, S1b); (3) the *vrn2-5* transcriptome has a large number of upregulated genes linked to cell-expansion. However, *vrn2-5* mutants also possess a larger root meristem (Labandera et al., 2021). Whether this is also the case in the *vrn2-5* SAM is yet to be established, but it is possible that VRN2-PRC2 also influences cell division in this tissue; we often observed faster leaf emergence in *vrn2-5* mutants, which is perhaps indicative of this. Interestingly, another N-degron pathway target, the transcription factor LITTLE ZIPPER 2 (ZPR2) was previously shown to coordinate leaf emergence from the hypoxic meristem in Arabidopsis (Weits et al., 2019), suggesting that multiple substrates of this proteolytic pathway may influence leaf organogenesis. However, the spatial disconnect between VRN2 localisation in the SAM and the ectopic expression of *pSAUR19::GUS* in the mature leaves of *vrn2-5* alludes to a meristem-derived or early developmental function for VRN2 that either persists through leaf maturation or is transduced across the leaf tissue. One possible mechanism is that targeting of *YUC8*, *YUC9* and *PIN1* by VRN2-PRC2 influences the synthesis or transport of bio-active auxins, which are known to potentiate PIF4 signalling (**Fig 5d**) (Franklin et al., 2011, Sun et al., 2012). A closer examination of auxin production, abundance, signalling, and transport in the *vrn2-5* mutant is therefore warranted.

Our data indicate that VRN2 is stable in the dividing regions of developing leaves but is destabilised via the N-degron pathway in the expansion region of mature leaves (**Figs 1b, 6d,e and S6**). This suggests that there may be a gradient of oxygen availability across the developing leaf, or that other signals impeding VRN2 turnover exist in the meristem and emerging leaves. For example, NO levels also feed into the control of N-degron substrates, including VRN2 (Gibbs et al., 2018, Hartman et al., 2019, Gibbs et al., 2014). Nonetheless, the question remains as to how proteolytic control of VRN2 across leaf development contributes to its light-dependent regulation of growth. As highlighted in our hypothetical model (**Fig. 6f**), stable VRN2 in the shoot meristem, emerging primordia and young leaves would ensure H3K27me3 is deposited at target loci. Since chromatin marks need to be replenished during mitosis, a continual pool of VRN2 in this region would lead to the maintenance of H3K27me3 at these loci as cells continually divide. As cells enter the expansion zone, H3K27me3 replenishment is no longer needed as cell division stops. This histone mark might then act as a scaffold/binding target for the PhyB/VIL1 module or other light-responsive regulators to introduce dynamic regulation in normoxic tissues. Indeed, it was previously shown that H3K27me3 levels in the *HAT4* REs remain constant across the dark-light cycle (Wang et al., 2024, Kim et al., 2021), indicating that H3K27me3 mark is stably imprinted prior to dynamic VIL1-mediated chromatin loop formation. Moreover, this spatiotemporally separated model is further reinforced by our observation that ectopic VRN2 accumulation does not lead to gain-of-function phenotypes – e.g., *prt6-1* mutant rosettes and hypocotyls are not significantly smaller than Col-0 plants (**Figs. 1e and 2d**), and that *pSAUR19::GUS* levels are not depleted in *prt6-1* relative to Col-0 (**Fig. 3a**).

Although VRN2-signal appears to be absent in mature leaves, it is possible that a small pool of stable VRN2 remains in these cells as part of the PRC2, for example due to shielding of its N-terminus. Under this mechanism, VRN2 might be maintained at PRC2 target genes to directly facilitate the binding of secondary regulators, whilst also maintaining H3K27me3 levels. Interestingly it was recently reported that emerging leaves are subject to a time-of-day dependent hypoxic cycle linked to starch availability during the night (Triozzi et al., 2024), which might also lead to transient stabilisation of VRN2 in areas of the plant that normally appear normoxic. A recent study also showed that the histone demethylase RELATIVE TO EARLY FLOWERING 6 (REF6), an antagonist of PRC2 that catalyses the removal of H3K27me3, also targets PIF4 regulated genes during thermomorphogenesis (He et al., 2022), suggesting that complex and dynamic chromatin modifications occur at these loci to tightly coordinate their expression.

To conclude, our work has identified a novel and developmentally encoded role for the polycomb protein VRN2 in restricting PIF4 activation of growth promoting genes in the light. This additional layer of regulation acts to ensure effective suppression of these genes in response to the environment and indicates that phytochrome-triggered degradation of PIFs alone is not sufficient to effectively switch off PIF targets. The epigenetic regulation of these targets provides new insight into the impact of H3K27me3 deposition in Arabidopsis development and provides increasing evidence that hypoxic gradients and dynamic chromatin remodelling both have significant roles to play in regulating environment-responsive growth.

## Materials and Methods

### Arabidopsis growth and transgenic lines

Arabidopsis seed were sown directly to compost (reduced peat Sinclair potting media, vermiculite, perlite, 4:2:1) or on ½ Murishage and Skoog (MS) medium with 1% agar. Plants were grown in a Weiss Technik fitotron SGC 120 growth chamber under long (16h) day conditions at 22°C (100 mol m^-2^•s^-1^). *vrn2-5, prt6-1, prt6-1vrn2-5, pif4-2* and *vil1-1* (also known as *vrn5-8*) were described previously (Gibbs et al., 2018, Kim et al., 2021, Leivar et al., 2008, Greb et al., 2007). Lines expressing *pSAUR19::GUS* (Spartz et al., 2012) were generated through crossing. The *pVRN2::VRN2-GUS/vrn2-5* and *pVRN2::VRN2-GUS/prt6-1* were previously reported (Gibbs et al., 2018). The *vrn2*/VRN2-YFP complementation line was generated by amplifying the genomic sequence of *VRN2,* including ∼2kb of promoter, by PCR ligating into pDONR221 and then mobilising into pGWB540 before transforming into *vrn2-5* mutant plants using the floral dip method. Homozygous transgenic plants were selected by growing seedlings on ½ MS media containing 15μg/ml Hygromycin B. Briefly, seed were sown on square plates and after stratification for 48 hours at 4°C were transferred to the light at 22°C for 4 hours before being wrapped in aluminium foil for a further 48 hours to promote skotomorphogensis. Aluminium foil was then removed, and plants were allowed to grow normally in the light for 7 days. Seedlings developing true leaves were selected for propagation and genotyped via PCR.

### Phenotyping assays

For rosette measurements, plants were grown in compost under long day (16h L: 8h D) conditions (100 mol m^-2^•s^-1^) for 3 weeks. Leaves were removed sequentially by developmental number and transferred to 1% agar plates before imaging. For hypocotyl length measurements, sterilised seed were sown on ½ MS medium and stratified at 4°C for 48 hours. Plates were transferred to the light and grown for 7 days before photographs were taken. Photos of etiolated seedlings were taken at the indicated timepoints. To evaluate growth at 28°C, seed were stratified on ½ MS plates and then grown for 7 days at either 22°C or 28°C. In all cases, images were analysed using ImageJ.

### Histochemical staining and leaf pavement cell size

Plants expressing either *pVRN2::VRN2-GUS* or *pSAUR19::GUS* were stained at timepoints/conditions indicated. Plant material was incubated at 37°C for 1 h - 8 h in the dark suspended in GUS staining solution (1% potassium ferricyanide, 1% potassium ferrocyanide, 100 mM PBS pH 7.0, 2 mM X-gluc, 0.1% v:v Triton X-100. Seedlings were de-stained in clearing fixative (3:1 ethanol:acetic acid, 1% v:v Tween®) overnight. Cleared samples were mounted in 60% glycerol and imaged using a GX optical microscope. For leaf cell size measurements, leaf 3 from representative 2-week-old Col-0 and *vrn2-5* plants were dissected and cleared using destaining fixative. Leaves were then incubated in Toluidine blue O solution (0.1% w:v toluidine blue, 0.1% v:v Triton X-100) at 37°C for 5 minutes, rinsed, and mounted in water. 2 images for 10 leaves of each genotype were captured from the left and right expansion zones of the midrib respectively using a GX optical microscope. Cell size and number were quantified using the Python tool LeafNet (Li et al., 2022).

### gDNA extraction for genotyping

Individual leaves of 3-week-old plants were harvested and frozen in liquid N_2_. Tissues were ground using a Qiagen Tissue Lyser II and gDNA was extracted using the Qiagen DNeasy® Plant Mini kit according to manufacturer instructions. Purified DNA was used as template for PCR with respective primers to genotype segregating transgenic plants.

### Western blotting

Proteins were extracted from whole Arabidopsis seedlings or dissected rosette leaves grown at 22°C under long day conditions (100 mol m^-2^•s^-1^). Tissues were ground in tubes using a Qiagen Tissue Lyser II and mixed with protein extraction buffer I (150 mM Tris-HCl pH7.5, 150 mM NaCl, 5 mM EDTA, 2 mM EGTA, 5% v:v glycerol, 0.1% SDS, 1% Triton X-100, 1x Complete Protease Inhibitor Cocktail (Roche)) at a ratio of 1:4. Protein quantification was performed using Bradford reagent (A6932, ITW Reagents) against a BSA standard curve. Total proteins were separated using SDS-PAGE and transferred to a PVDF membrane. PageRuler™ Plus prestained protein ladder (ThermoFisher) was used for reference of protein size. After transfer, membranes were blocked in 5% milk in TBS supplemented with 0.1% Tween®-20 for at least an hour. Membranes were probed overnight with antibodies in 5% milk in TBS supplemented with 0.1% Tween®-20; anti-GFP (1:1,000, Sigma-Aldrich, SAB4301138), anti-GUS (1:2000, Sigma-Aldrich, G5420), anti-PIF4 (1:1000, Agrisera, AS16 3955), anti-rabbit IgG conjugated to HRP (1:5000, Santa Cruz; sc-2004), anti-mouse IgG conjugated to HRP (1:5000, Santa Cruz; sc-358914), anti-goat IgG conjugated to HRP (1:10,000, Promega, PA1 28664), anti-H3 (1:2500, Abcam; ab1791), anti-H3K27me3 (1:1000, Millipore; 17-622).

### qRT-PCR

Whole seedlings were flash frozen at indicated timepoints for mRNA extraction. Tissues were ground in tubes using a Qiagen Tissue Lyser II and mRNA was extracted using RNeasy Plant Mini kit (Qiagen) according to manufacture instructions. mRNA was on-column DNase treated with DNAse Set (Qiagen). mRNA concentration was quantified using a NanoDrop spectrophotometer and 1 μg of cDNA was synthesised using the qScript cDNA Supermix kit. Quantitative real-time PCR was performed using a QuantStudio™ 5 Real-Time PCR System with Brilliant II low ROX Sybr Green (Agilent). Reactions were performed with at least 2 technical replicates and 3 biological replicates were performed for each experiment. Gene expression is expressed as 2^-ΔCT^ relative to two housekeeping genes. Primers for genes tested are shown in the key resources table.

### RNA-sequencing and analysis

For seedling RNA sequencing, plants were grown for 10 days on vertical ½ MS plates at 22°C under long day conditions (100 mol m^-2^•s^-1^) before being flash frozen in liquid N_2_ (3 biological replicates). Seedlings were ground in liquid N_2_ using a pestle and mortar and mRNA was extracted from 100 mg of leaf powder using the RNeasy Plant Mini kit (Qiagen) according to manufacture instructions. mRNA was on-column DNase treated with DNAse Set (Qiagen). Library preparation was performed by Glasgow Polyomics using the Illumina TruSeq mRNA kit (polyA selection). Paired end sequences of length 75 bp were generated using the HiSeq 4000 platform. Adapter ligated mRNA was trimmed and filtered using fastp with settings -l 50 and -q 20. Reads were aligned to the Arabidopsis genome (TAIR10) using bowtie and analysed using Kallisto^3^ before differential gene expression analysis was performed using the R package DEseq2^4^. Genes in *vrn2-5* or *prt6-1vrn2-5* were considered upregulated if they had an average log_2_ fold change of at least 0.58 across 3 biological replicates, corresponding to an upregulation of at least 1.5 relative to Col-0 and *prt6-1*, respectively.

For rosette RNA sequencing, plants were grown for 3 weeks at 22°C under long day conditions (100 mol m^-2^•s^-1^) in compost mix (reduced peat Sinclair potting media, vermiculite, perlite, 4:2:1) before being flash frozen in liquid N_2_. 4 independent biological replicates were performed, each of which consisted of tissue from a pool of 5 plants per genotype. Plants were ground in liquid N_2_ using a pestle and mortar and mRNA was extracted from 100 mg of leaf powder using the RNeasy Plant Mini kit (Qiagen) according to manufacture instructions. mRNA was on-column DNase treated with DNAse Set (Qiagen). Library preparation was performed by Novogene, UK. At least 20 million paired end sequences of length 150 bp per sample, per replicate, were generated using the Novoseq 6000 sequencing platform. Adapter removal, quality control, alignment, read counting and differential gene expression analysis were performed using HISAT2 (Zhang et al., 2021). Differential gene expression analysis was performed in R studio using the EdgeR (Robinson et al., 2010) and DESeq2 (Love et al., 2014) packages. Genes in *vrn2-5* were considered upregulated if they had an average log_2_ fold change of at least 0.58, corresponding to an upregulation of at least 1.5 relative to Col-0 plants. Gene ontology enrichment analysis was performed using the R package TopGO^(5)^. Venn diagrams were created using Venny 2.0 (https://bioinfogp.cnb.csic.es/tools/venny/index2.0.2.html).

### ChIP sequencing

For VRN2-YFP protein ChIP, 6g of 13-day old seedlings grown on ½ MS plates at 22°C under long day conditions (100 mol m^-2^•s^-1^) were harvested into 3 miracloth bags and washed in SDW. Seedlings were crosslinked by vacuum infiltration for 15 minutes in PBS supplemented with 1% formaldehyde. The crosslinking reaction was quenched by adding 2M glycine to a final concentration of 125 mM and applying a vacuum for a further 5 minutes. Seedlings were then washed in SDW 4 times before being patted dry to remove excess moisture and flash frozen in liquid nitrogen. Tissues were ground in liquid nitrogen using a pestle and mortar for at least 10 minutes. Nuclei were isolated using 70 ml HONDA buffer (20 mM HEPES pH 7.4, 440 mM sucrose, 1.25% w:v Ficoll, 2.5% w:v Dextran-T40, 10mM MgCl_2_, 0.5% v:v Triton X-100, 5 mM DTT, 1x Complete Protease Inhibitor Cocktail (Roche), 25 μM MG-132 (Sigma Aldrich)). After addition of HONDA buffer, samples were gently rotated at 4°C for 15 minutes. Extracts were centrifuged at 3,500 g for 30 minutes at 4°C to pellet nuclei. The supernatant was then removed and nuclei were gently resuspended in 1.5 ml of HONDA buffer and split into 3 x 1.5 ml DNA LoBind tubes (Eppendorf). Samples were centrifugated at 2,000 g for 10 minutes to pellet nuclei and the supernatant was removed. This washing process was repeated 2 times for each sample. Nuclei were resuspended in nuclear lysis buffer (50 mM Tris-HCl pH 8.0, 10 mM EDTA, 1% v:v SDS, 1x Complete Protease Inhibitor Cocktail (Roche), 25 μM MG-132 (Sigma Aldrich)) and sonicated using a Bioruptor® Plus (Diagenode) for 3 x 5 minutes on low setting to lyse nuclei and shear chromatin to generate fragments between 200 and 800 bp, centring around 500 bp. Samples were then centrifuged at 14,500 g for 5 minutes at 4°C to pellet cell debris. Chromatin extracts were transferred to 15 ml LoBind tubes and diluted 1:10 in ChIP dilution buffer (15 mM Tris-HCl pH 8, 1.2 mM EDTA, 1.1% v:v Triton X-100), and precleared with 50 μl of Protein-A agarose (ab193254) by gentle rotation at 4°C. For each biological sample, 3 IPs were performed independently from the equivalent of 2 g of chromatin and pooled during DNA cleanup. Prior to immunoprecipitation 150 μl of chromatin extract was taken as input control. 4 μg of anti-GFP (ab290, Abcam) antibody was incubated with chromatin extracts for 3 hours and gently rotated at 4°C. VRN2-antibody complexes were immunoprecipitated overnight using 50 μl of Protein-A agarose. Beads were sequentially washed twice each by rotating for 5 minutes at 4°C with low salt (150 mM NaCl, 0.1% v:v SDS, 1% v:v Triton X-100, 4mM EDTA, 20 mM Tris-HCl pH 8.0), high salt (500 mM NaCl, 0.1% v:v SDS, 1% v:v Triton X-100, 4mM EDTA, 20 mM Tris-HCl pH 8.0) and TE (2 mM EDTA, 10 mM Tris-HCl pH 8.0) buffers. DNA-protein complexes were eluted twice with 200 μl ChIP elution buffer (1% v:v SDS, 100 mM NaHCO_3_) by incubating for 15 minutes at 65°C. Samples were reverse crosslinked overnight by incubating at 65°C with 200 mM NaCl before treatment with Proteinase K for 1 hour at 65°C. ChIPed DNA was purified using phenol, chloroform, isoamyl alcohol 25:24:1 (v/v/v) (Sigma) followed by precipitation at −20°C overnight in 100% EtOH with 90 mM NaOAc and 1μl GlycoBlue™ Coprecipitant (ThermoFisher). Precipitated DNA was centrifuged at 21,000 g for 1 hour at 4°C and washed twice with 70% EtOH. The DNA pellet was resuspended in 50 ul of TE buffer and cleaned up further using the ChIP DNA Clean and Concentrator™ kit according to manufacture instructions. Library preparation and sequencing was performed by Glasgow Polyomics. A minimum of 1 ng of ChIPed DNA was processed using the NEBNext® Ultra™II DNA Library Prep kit according to manufacturer instructions. Adapter ligated DNA was amplified with 10 rounds of PCR and sequenced 2×100bp using the Illumina NextSeq2000 sequencing system.

For histone H3K27me3 ChIP sequencing 1 g of 11-day old seedlings grown in ½ MS plates were harvested in 50 mL falcon tubes and washed twice in SDW. Seedlings were crosslinked by vacuum infiltration in PBS containing 1% (v/v) formaldehyde 15 min. Crosslinking reaction was quenched by the addition of a final concentration of 125 mM glycine and applying further vacuum for 5 min. Crosslinked seedlings were rinsed 4 times with SDW and patted dried and flash frozen. Tissues were ground with mortor and pestle and resuspended in 30 mL extraction buffer 1 (10 mM Tris-HCl, pH 8.0, 5 mM βmercaptoethanol, 0.4 M sucrose, protease inhibitor cocktail (cOmplete, Roche), vortexed, incubated for 5 min and filtered through 2 layers of Miracloth. Filtered solution was centrifuge at 2880 g for 20 min at 4°C and the pellet resuspended in 1 mL extraction Buffer 2 (10 mM Tris-HCl, pH 8.0, 0.25 M sucrose, 10 mM MgCl2, 1% (v/v) Triton X-100, 5 mM β-mercaptoethanol, 1x protease inhibitor cocktail (cOmplete, Roche)). Solution was transferred to a new 1.5 mL Eppendorf tube and centrifuged at 12000 xg for 10 min at 4°C. Pellet was resuspended in 300 μL extraction buffer 3 (10 mM Tris-HCl, pH 8.0, 1.7 M sucrose, 0.15% (v/v) Triton X-100, 2 mM MgCl2, 5 mM β-mercaptoethanol, 1x protease inhibitor cocktail (cOmplete, Roche)). The resuspended pellet was overlayed onto 900 μL extraction buffer 3 and centrifuge at 16000 xg for 1h at 4°C. The chromatin pellet was resuspended in 320 μL nuclei lysis buffer (50 mM Tris-HCl, pH 8.0, 10 mM EDTA, 1% (w/v) SDS, 1x protease inhibitor cocktail (cOmplete, Roche)) and transferred to a DNA LoBind tube. An aliquot of 10 μL (unsheared chromatin) was saved for later quality check. Chromatin sonicated 3 times for 5 min (30 sec on/off intervals) in a Bioruptor® Plus (Diagenode) at the low setting to generate DNA fragments in the range of 200-800 bp. Sheared chromatin was centrifuged at 16000 xg for 5 min and supernatant kept. An aliquot of 10 μL (sheared chromatin) was saved for later quality check. Dynabeads Protein A magnetic beads (15 μL per sample) were washed three times in ChIP dilution buffer (1.1% (v/v) Triton X-100, 1.2 mM EDTA, 16.7 mM TrisHCl, pH 8.0, 167 mM NaCl, 1x protease inhibitor cocktail (cOmplete, Roche). Antibodies were added to the beads in 50 μL ChIP dilution buffer (4 μg anti-H3K27me3, Millipore 17-622) and incubated for 1 h at 4°C. The beads were washed 3 times with ChIP dilution buffer. 10% of the chromatin solution was kept as input sample. The remaining chromatin was diluted 10 times in ChIP dilution buffer and 900 μL were added to each antibody bound beads and incubated by rotation overnight at 4°C. Each sample was washed 2x 5min with 1 mL Low salt wash buffer (150 mM NaCl, 0.1% (w/v) SDS, 1% (v/v) Triton X-100, 2 mM EDTA, 20 mM Tris-HCl, pH 8.0), and 1x 5min with 1 mL high salt wash buffer (500 mM NaCl, 0.1% (w/v) SDS, 1% (v/v) Triton X-100, 2 mM EDTA, 20 mM Tris-HCl, pH 8.0). 1 mL of Litium Chloride wash (0.25 M LiCl, 1% (v/v) NP-40, 1% (w/v) sodium deoxycholate, 1 mM EDTA, 10 mM Tris-HCl, pH 8.0) was added and incubated for 5 min followed by the addition of 1 mL of TE buffer (10 mM Tris-HCl, pH 8.0, 1 mM EDTA). Beads were finally washed with 1 mL TE buffer for 5 min. The immune complexes were eluted from the beads with 2x 250 μL 1% SDS, 0.1 M NaHCO_3_ for 20 min at 65°C. Reverse crosslinking was performed by the addition of a final concentration of 0.3 M NaCl and incubated overnight at 65°C. DNA was purified with 500 μL phenol, chloroform, isoamyl alcohol 25:24:1 (v/v/v) (Sigma) and precipitated with 1.35 mL ice cold 100% EtOH, 50 μL NaAc 3 M, pH 5.2, 2 μL glycoblue, incubated 72h at −80°C. Samples were centrifuged at maximum speed for 5 min and a pellets were washed 2 times with 75% EtOH followed by air drying.

### ChIP sequencing Analysis

For VRN2-YFP protein ChIP, adapters ligated to raw paired-end reads were trimmed and filtered using FastP^8^ with minimum length set to 50 bp and minimum q score of 20. Trimmed reads were aligned to the TAIR10 assembly of the Arabidopsis genome using bwa-mem of the Barrows-Wheeler Alignment Tool (Li and Durbin, 2009)with default parameters. Alignments were subject to an additional filter in which reads were removed if they were unmapped or had a mapping quality score of <30. Prior to peak calling, a blacklist of highly duplicated regions was removed from all alignment files (Klasfeld et al., 2022). MACS2 (Zhang et al., 2008) was used to call enriched regions with the settings -g 101274395 -f BAMPE --nomodel –keep-dup auto -q 0.05. Consensus peaks for VRN2-YFP were considered if they were observed to have an FDR of < 0.05 and an enrichment of at least 1.5 fold in at least 2 independent biological replicates. The closest gene to each peak was annotated using the annotatePeak function in the R package ChIPSeeker (Yu et al., 2015)with priority set to promoters, 5UTR, 3UTR, exon, intron, downstream and intergenic regions. For differential H3K27me3 ChIP, adapters ligated to raw single-end reads were trimmed and filtered using FastP (Chen et al., 2018) with minimum length set to 50 bp and minimum q score of 20. Trimmed reads were aligned to the TAIR10 assembly of the Arabidopsis genome using bwa-mem of the Burrows-Wheeler Alignment Tool (Li and Durbin, 2009) with default parameters. Alignments were subject to an additional filter in which reads were removed if they were unmapped or had a mapping quality score of <30. Prior to peak calling, a blacklist of highly duplicated regions was removed from all alignment files(Klasfeld et al., 2022). SICER2 (Xu et al., 2014) was used to perform differential peak calling with the sicer_df function, with settings -egf 0.95 –significant_reads. Consensus hypomethylated peaks in *vrn2-5* were considered if they were observed to have an FDR of < 0.05 and a relative fold change difference of at least 80% of WT in at least 2 independent biological replicates. The closest gene to each peak was annotated using the annotatePeak function in the R package ChIPSeeker (Yu et al., 2015) with priority set to promoters, 5UTR, 3UTR, exon, intron, downstream and intergenic regions. Where peaks mapped to multiple genes, only the closest was considered. To visualise data, bigwig files were created using the DeepTools (Ramirez et al., 2016)function bamCoverage to bin reads into 20 bp windows which were normalised by reads per genome coverage (RPGC). Gene specific enrichment was evaluated using IGV.

### ChIP qPCR

The ChIP PCR protocol has previously been described (Franco-Echevarria et al., 2023). Samples were handled exactly as for the VRN2-YFP ChIP sequencing, with some differences. Isolated nuclei were resuspended in RIPA buffer (1% NP-40 (v:v), 0.1% sodium deoxycholate (v:v), 0.1% SDS (v:v), 1x Complete Protease Inhibitor Cocktail (Roche), 25 μM MG-132 (Sigma Aldrich) in PBS). Nuclei were lysed by sonication on high setting for 5 minutes on a cycle of 30s on, 30s off, twice. For VRN2-YFP samples, chromatin was sheared for 15 minutes by treatment with the endonuclease benzonase; for H3K27me3 ChIP, nuclei were lysed by sonication and DNA fragmentation was achieved using 3 cycles of 30s on 30s off for 5 minutes on low setting. For VRN2-YFP, the IP step was performed overnight using 4 μg of anti-GFP (ab290, Abcam) and Protein-A agarose (ab193254) which had been blocked with salmon sperm DNA (10 mg/ml, ThermoFisher). For H3K27me3, the IP was performed overnight with 3 μg of anti-H3K27me3 (Millipore, 17-622) or anti-H3 (Abcam, ab1791) respectively with Protein A magnetic Dynabeads. For each qPCR, four technical repeats were performed from a minimum of 2 independent biological replicates. Enrichment values were calculated by normalising the % input to the enrichment of *5SrDNA* which served as a negative control.

### Confocal microscopy

VRN2-YFP seed were stratified in the dark at 4°C for 48 hours, and then grown vertically ½ MS agar plates for 4 days. Seedlings were then mounted in 60% glycerol and imaged using a Zeiss LSM 710 confocal microscope.

## Author contributions

D.J.G and R.O designed the study and its experiments. R.O, A-M.L, A.J.R, A.K. O.A, M.A.S, C.M, T.X, S.H, E.K and D.J.G conducted experiments. R.O, A-M.L, S.H, E.K and D.J.G analysed data. S.H, E.K and D.J.G secured funding for the work. D.J.G and R.O wrote the paper with input from all authors.

## Acknowledgements.

We thank William Gray (University of Minnesota, USA) for sharing the *pSAUR19::GUS* reporter line, Daan Weits (Utrecht University, Netherlands) for sharing the *pHRPE::GUS* reporter line, Hannes van Haeren (VIB, Ghent, Belgium) for advice with rosette quantification, Jie Song (Imperial Gollege London, UK) and Anna Schulten and Robert Maple (John Innes Centre, UK) for advice on ChIP-seq, and Rashmi Sasidharan (Utrecht University, Netherlands) for early discussions on red/far-red light signalling. D.J.G. was supported by a European Research Council (ERC) Starting grant (GasPlant-715441) and Biotechnology and Biological Sciences Research Council (BBSRC) grant BB/V008587/1. Work in the lab of E.K was supported by BB/X009653/1. S.H was supported by the Netherlands Organization for Scientific Research (019.201EN.004) and the Deutsche Forschungsgemeinschaft (DFG, German Research Foundation) under Germany’s Excellence Strategy (CIBSS-EXC-2189-Project ID 390939984). M.S. was supported by a BBSRC funded MIBTP-DTP studentship.

---

**Figure S1.**
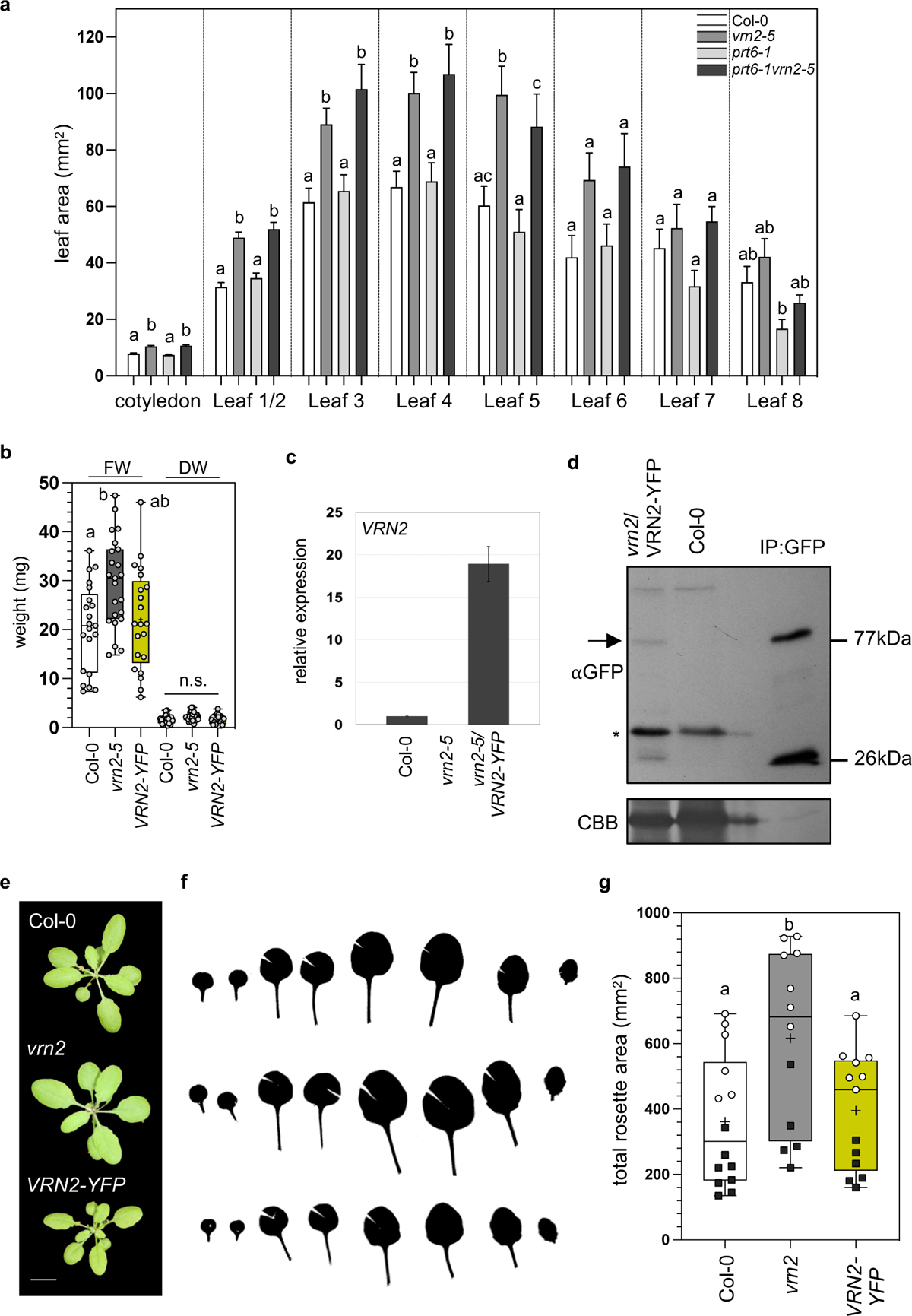
(**a**) Quantification of individual leaves of Col-0, *vrn2-5*, *prt6-1* and *prt6-1vrn2-5*. Bars represent the average value from at least 20 leaves from 4 biological replicates. Error bars represent the standard error of the mean. Letters indicate statistically significant differences between genotypes for each numbered leaf and were calculated by 1-way ANOVA followed by Tukey’s test. (**b**) Fresh (FW) and dry (DW) measurements of 4-week-old rosettes of Col-0, *vrn2-5* and *vrn2-5/VRN2-YFP*. Statistical differences within conditions were calculated by one way ANOVA followed by Tukey’s test. (**c**) qRT-PCR quantifying *VRN2* expression in light grown seedlings of Col-0, *vrn2-5* and *vrn2-5*/*VRN2-YFP*. Data were normalised to 2 housekeeping genes and are presented relative to Col-0. Each bar represents the average of data from 2 biological replicates, error bars represent the standard error of the mean. (**d**) Western blot showing VRN2-YFP protein abundance (arrow, right) in *vrn2-5/VRN2-YFP* plants. Numbers indicate protein size (kDa). Col-0 was used as a negative control, whilst the GFP:IP was used to demonstrate the transcribed protein can be immunoprecipitated. A non-specific band (*) is present in input samples, and the band at 26 kDa in the IP lane corresponds to free YFP. (**e**) Representative rosettes of Col-0, *vrn2-5* and *vrn2-5*/VRN2-YFP. Note that the Col-0 and *vrn2* rosettes are duplicated from main Fig. 1c. (**f**) Representative silhouettes of dissected leaves assembled in decreasing developmental age from left to right for Col-0 (top), *vrn2-5* (middle) and *vrn2-5*/VRN2-YFP (bottom). (**g**) Box and whisker plot showing total rosette area for Col-0, *vrn2-5* and *vrn2-5*/VRN2-YFP. Data points represent individual plants from 2 biological replicates (denoted by circles or squares). These data are also included alongside extra reps for Col-0, *vrn2-5*, *prt6-1* and *prt6-1 vrn2-5* in main Fig. 1e. Letters indicate statistically significant differences calculated by 1-way ANOVA followed by Tukey’s test.

**Figure S2.**
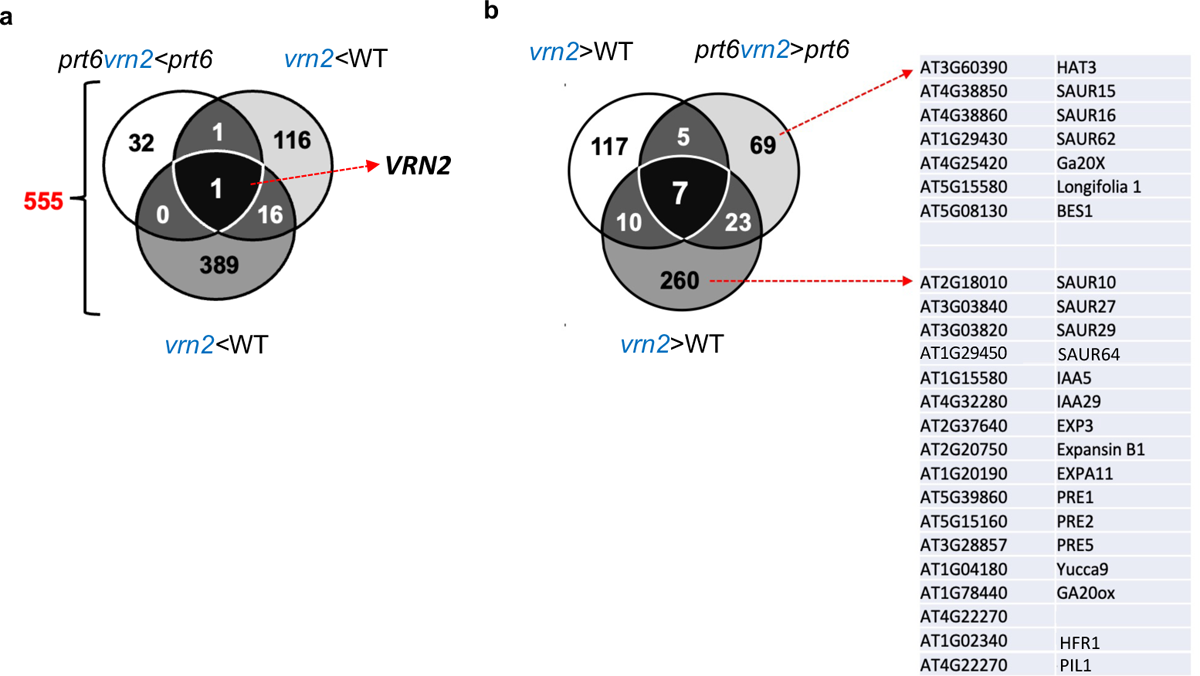
(**a**) Venn diagram showing the overlap of downregulated genes in *vrn2-5* vs Col-0 at the seedling and rosette stage, and *prt6-1 vrn2-5* vs *prt6-1* at the seedling stage. (**b**) Venn diagram as in main Fig. 2a – callouts list genes which were represented by the GO terms ‘*response to growth*’.

**Figure S3.**
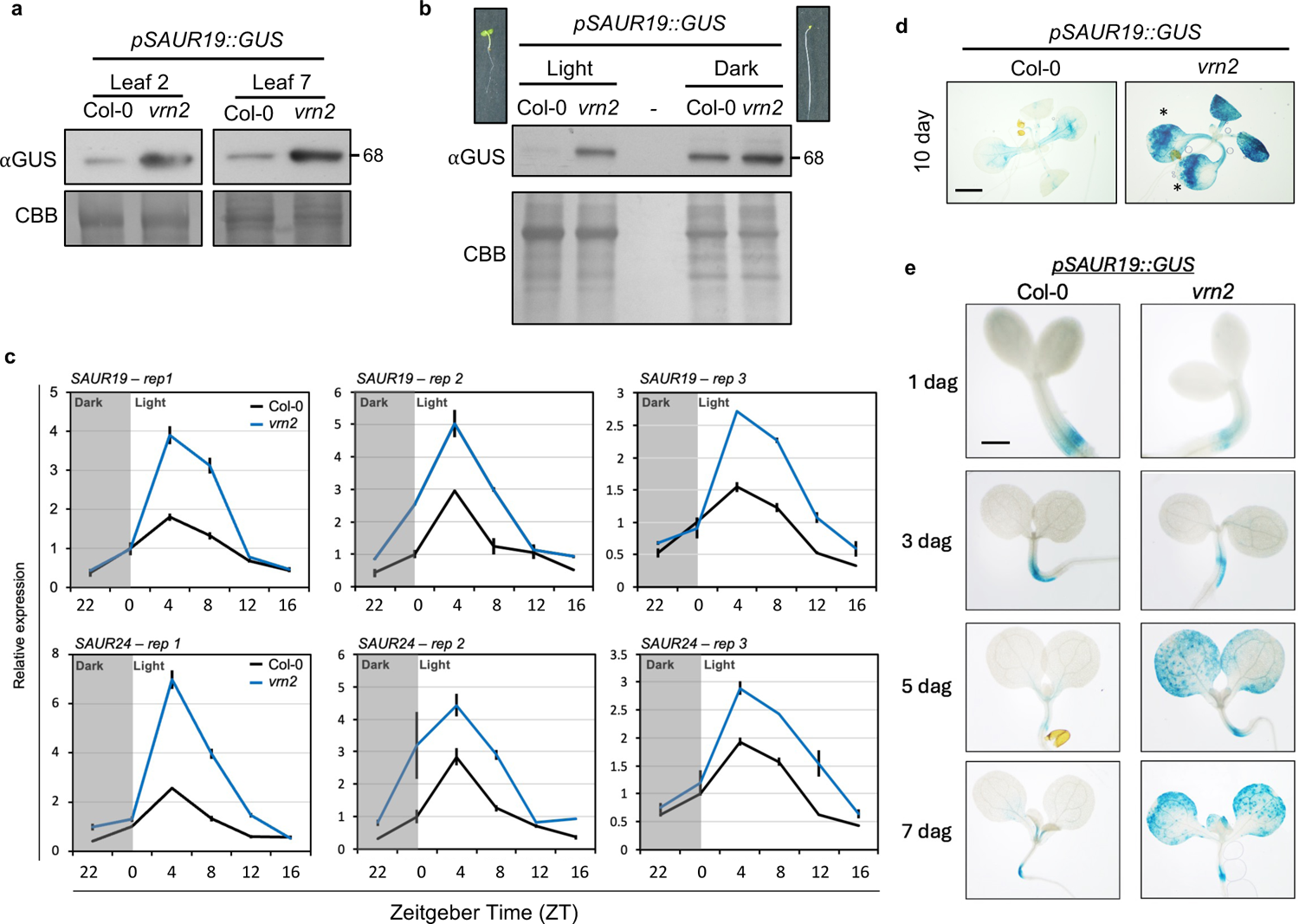
(**a**) anti-GUS Western blot of *pSAUR19::GUS* in leaf 2 and leaf 7 of 4-week-old Col-0 and *vrn2-5* plants. Coomassie brilliant blue (CBB) staining was used to quantify equal loading between samples. (**b**) anti-GUS Western blot of *pSAUR19::GUS* in 4-day-old light vs dark grown Col-0 and *vrn2-5* seedlings. Coomassie brilliant blue (CBB) staining was used to quantify equal loading between samples. (**c**) qRT-PCR quantifying *SAUR19* and *SAUR24* expression in Col-0 and *vrn2-5* at the indicated timepoints displayed in Zeitgeber (ZT) time across 3 independent biological replicates (Rep 1 is also shown in main Fig. 3). Data were normalised to 2 housekeeping genes and are presented relative to Col-0 at ZT 0. Each point represents the mean of 2 technical repeats; error bars represent the standard error of the mean. (**d**) Histochemical staining of representative 10-day old seedlings of Col-0 and *vrn2-5* plants expressing *pSAUR19::GUS*. Scale bar = 1mm. Note the pattern of ectopic GUS expression in *vrn2-5*, which corresponds to the cell expansion zone on the outer edge of true leaves (denoted with asterisks). (**e**) Histochemical staining of representative Col-0 and *vrn2-5* seedlings expressing *pSAUR19::GUS* at 1, 3, 5 and 7 days after germination (dag).

**Figure S4.**
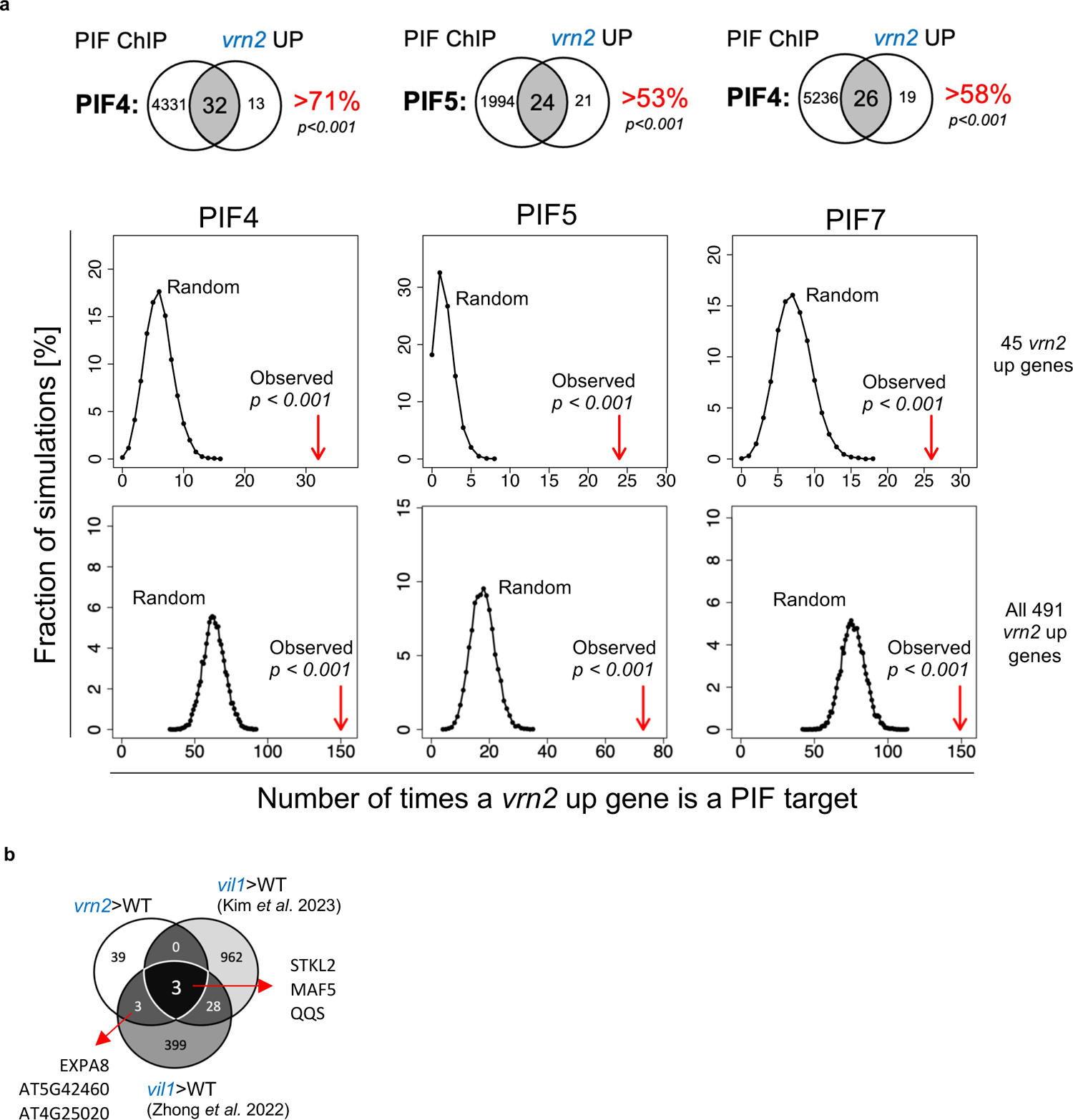
(**a**) Venn diagrams describing the overlap of the 45 *vrn2-5* up genes with reported PIF4, PIF5 and PIF7 binding targets. Graphs show the simulated overlap vs the observed overlap (Obs., red arrow). Data are plotted as a frequency distribution across 10,000 random simulations. Red arrows to the right of this distribution indicate the observed overlap is not likely to have been observed by chance. (**b**) Venn diagrams describing the overlap of the 45 *vrn2* up genes with the genes previously reported as upregulated in 1- and 14-day old *vil1* seedlings.

**Figure S5.**
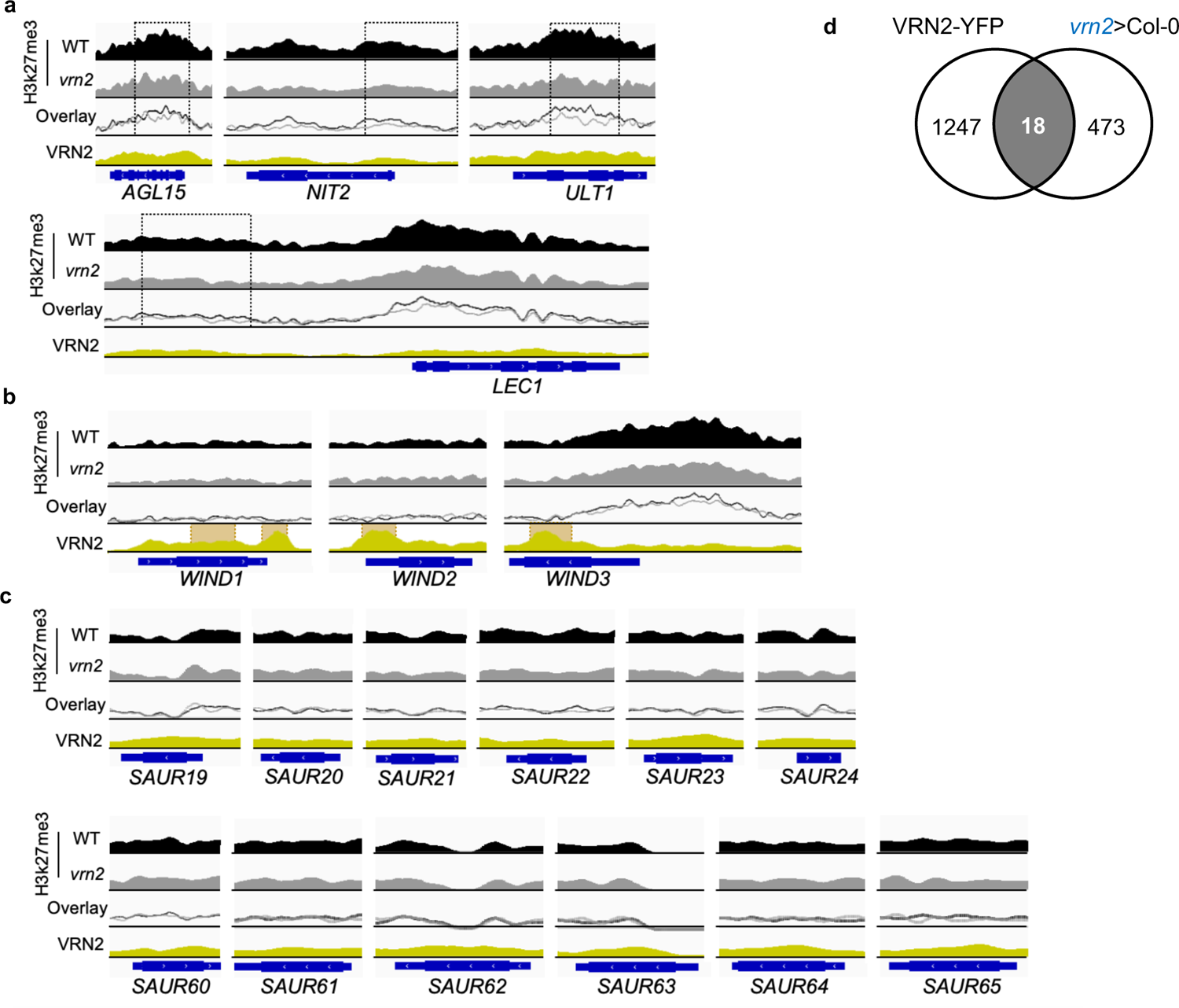
(**a**) IGV plots showing alignment data for previously reported PRC2 targets *AGL15*, *NIT2*, *ULT1* and *LEC1*. Black and grey tracks represent Col-0 and *vrn2-5* H3K27me3 coverage respectively; ‘overlay’ represents these same tracks mapped over one another. Black dotted boxes indicate regions which were identified as significantly depleted for H3K27me3 in *vrn2-5*. The yellow track represents VRN2-YFP coverage across the same genomic loci; orange boxes indicated regions which were identified as significantly enriched by VRN2-YFP. Data show coverage across the combination of 3 biological replicates which were normalised by reads per genome coverage. In each graph, blue tracks indicate the position and direction of the genomic sequence for each gene. (**b**) As in (a) for the previously reported VRN2 targets *WIND1*, *WIND2* and *WIND3*. (**c**) As in (a) for the *SAUR19-24* and *SAUR60-65* gene families, respectively. Note that no regions were identified as bound by VRN2-YFP or having depleted levels of H3K27me3 in *vrn2-5*. (**d**) Venn diagram showing the overlap between VRN2-YFP targets and the 491 *vrn2-5* UP DEGs.

**Figure S6.**
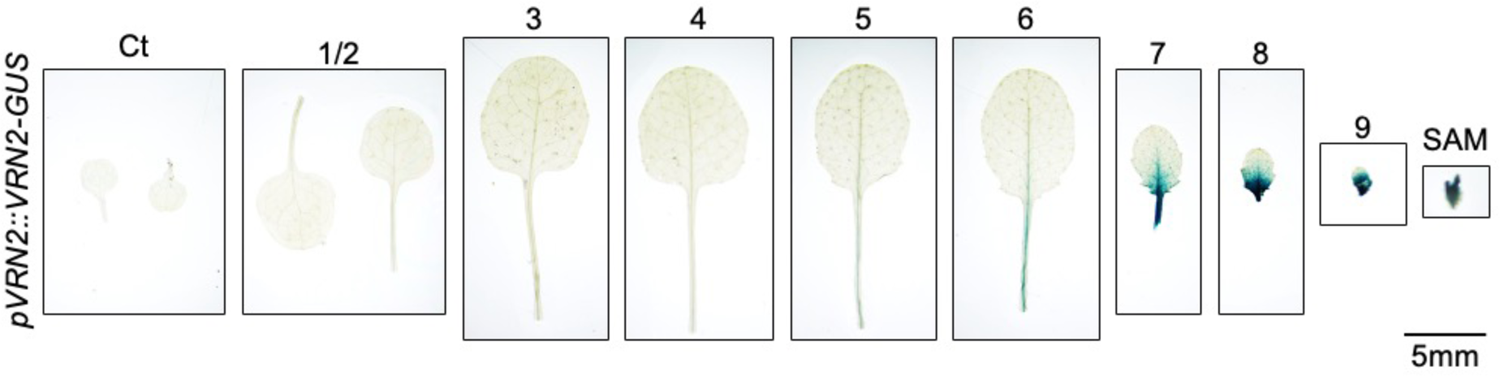
Representative histochemical staining of all leaves and tissues from the plant in Fig. 6d and 6e. Leaves are arranged in developmental order from oldest (left) to youngest (right), as well as the shoot apical meristem (SAM) on the right. Scale bar = 5 mm.

